# BAGEL-CAR: Reflections on the *Bits to Binders* Competition

**DOI:** 10.64898/2026.09.02.748856

**Authors:** Jakub Lála, Stefano Angioletti-Uberti

## Abstract

The antigen-binding segment of chimeric antigen receptors (CARs) in CAR-T therapy has emerged as a compelling application of *de novo* AI protein design. In the *Bits to Binders* competition, our group submitted 414 BAGEL-CAR designs for CD20-directed CAR binding segments, 38.4% of which were statistically enriched in a pooled CAR-T proliferation screen, the highest among all teams and more than double the next-best. Strikingly, aŁer detailing the three design strategies employed, we show that two of them carry the entire success of our campaign, with both of them outcompeting all the other teams’ aggregate results. Crucially, during the competition, we did not refold any of the binders aŁer backbone painting. Had we applied this typical validation step with ESMFold, all experimentally enriched designs would have been filtered out. Our findings show that each filtering step should be stress-tested per biomolecular system and intermediate outputs of design pipelines should be assayed regardless of ‘common practice’. Additionally, we appear to be the only team that designed against ESMFold’s predicted CD20 dimer geometry, which diverges substantially from the experimentally resolved `6Y97` structure, raising, but not resolving, the possibility that this alternative target geometry contributed to our high hit rate. We hope these reflections help inform the future of effective CAR-T and binder design campaigns.

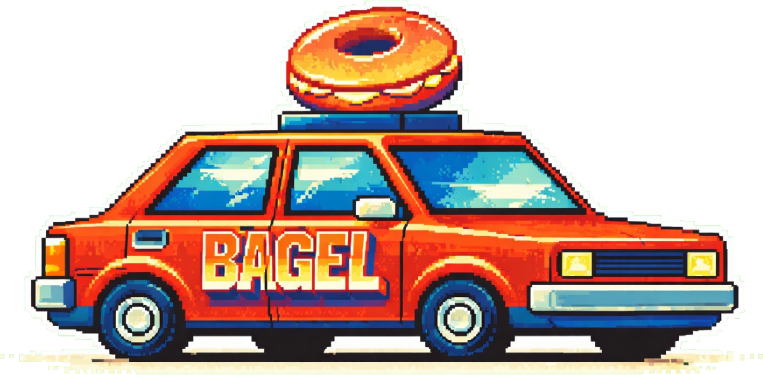

## 1 Introduction

CAR-T therapy involves designing a chimeric antigen receptor (CAR), and introducing it into an extracted T cell *ex vivo*. Such an engineered CAR-T cell can then be infused into a patient, with the goal of removing malignant cells in cancer [1], or autoreactive B cells in autoimmune disease [2]. Although AI protein design could in principle address the full CAR construct *de novo*, progress on protein binders [3], [4], [5] singles out the binding segment as the tractable entry point by introducing such binders into a predefined construct with validated, fixed transmembrane and signaling domains. The question then remains as to how one designs a binder engaging the target receptor on the surface of diseased cells, considering the fact that this binder needs to be physically attached to a longer CAR construct that should remain functional in terms of signaling.

To that end, Kosonocky et al. [6] recently released the results from the *Bits to Binders* competition, showing the potential of *in silico* protein design methods to produce such CARs. In the competition, 28 teams submitted 12,000 designed 80-amino-acid binders against human CD20. These were tested in a pooled CAR-T proliferation assay, in which each designed binder also served as its own DNA barcode for its CAR-T cell, and enrichment was measured as differential expansion upon co-culture with CD20+ cancer cells vs. no-target controls. The top 10 enriched CARs were then taken forward into individual assays, including proliferation, expansion, cytokine production, cytotoxicity, and binding measurements.

The competition revealed that current computational protein design tools can produce functional CAR binding domains. Out of 12,000 total designs, 707 were significantly enriched, with team hit rates of 0.6%–38.4%. Notably, many commonly used *in silico* filtering metrics from protein design workflows could not predict the experimental outcome of the CAR-T proliferation in the pooled assay, calling into question the reliability of these computational filters for predicting binding and functional outcomes.

To put this in context, consider the typical workflow pioneered by the Baker Lab and validated in numerous experiments [7], [8], [9]. First, a *generation* process proposes a backbone (potentially co-proposed with a sequence) against a chosen epitope, oŁen with diffusion-based models [10], [11], [12] or so-called hallucination algorithms [13], [14]. Second, this backbone is assigned a specific amino acid sequence using an inverse folding algorithm [15], [16], also referred to as *inpainting*. While such algorithms provide the sequence most likely to fold into that input backbone, this does not mean that the specified backbone is the most likely or energetically favorable state for that sequence; it means only that alternative sequences are less likely to adopt that specific fold. This consideration motivates a third and final step, *refolding validation*, in which the binder is folded together with its target to check that the predicted structure remains close to the initial design and that confidence metrics characterizing the binder-target interface, such as ipSAE [17], exceed a threshold consistent with a binding interaction.

This last step, in principle, provides some form of self-consistency to the whole pipeline by seeking consensus across conceptually orthogonal models: diffusion-based backbone generators, graph-based autoregressive sequence painting, and structure prediction models. More importantly, if one could fully trust the validation folding algorithm, details of the preceding backbone and sequence generation should be almost irrelevant. In other words, it does not matter how one arrives at a specific binder sequence as long as the validation algorithm, or more precisely our interpretation of its output, predicts that a binding interface is formed. From here on, we refer to the entire generative pipeline as the “generation → refolding validation” pipeline. Where appropriate, we expand the generation stage into the traditional “backbone generation → fixed-backbone sequence design” steps.

Therefore, under this definition, our previously described method BAGEL [18] collapses these two steps into a single sampling procedure in which the generation step samples directly from the final validation oracle via Monte Carlo (MC) optimization. This can also be viewed as a non-gradient-based hallucination approach [19]. Using this method, together with further filtering steps, we nominated 414 BAGEL-derived designs for wet-lab validation as part of the competition under the team name *Nucleate UK London*. We refer to these sequences as BAGEL-CAR designs. Our team achieved the highest pooled proliferation hit rate, 38.4%, more than twice that of the second-best team (Binding Illini at 14.8%, using Chroma [12]). One of our designs was further validated by significant specific lysis and confirmed binding with *K_D_* ≈ 643 nM. Genuinely surprised by the extent of our success, and motivated to elucidate our design approach and rationale, we write this short follow-up piece. To avoid reiterating most of Kosonocky et al., we summarize the competition results in Figure 1 and refer the reader to the original preprint [6].

**Figure 1:**
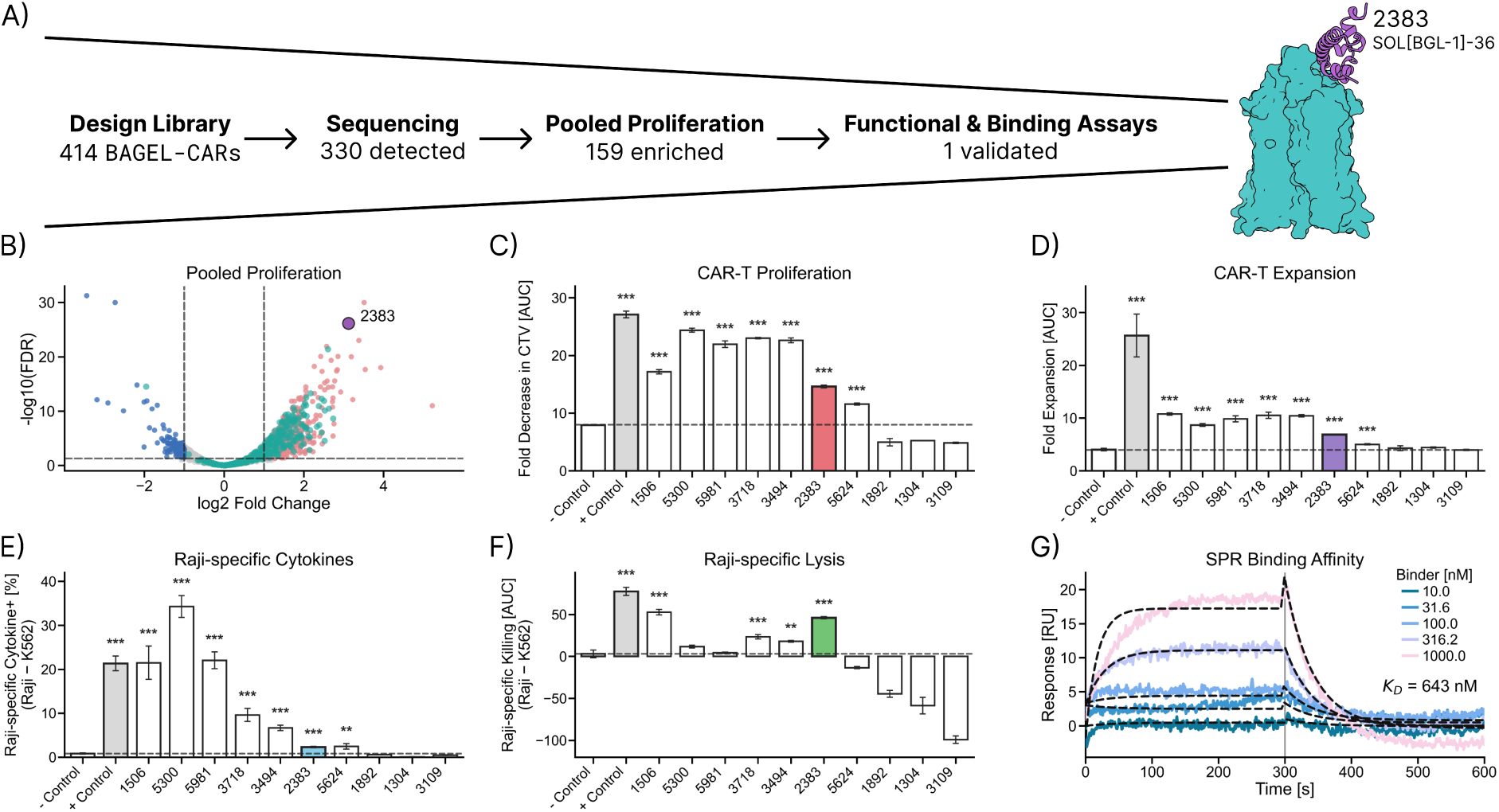
Results for the BAGEL-CAR Designs in the *Bits to Binders* Competition. Competition assay data for all designs, with BAGEL-CAR designs highlighted. For experimental details, see Kosonocky et al. [6]. **A)** BAGEL-CAR screening funnel: 414 submitted, 330 sequencing-detected, 159 enriched in the pooled CAR-T proliferation assay, 1 binder (`SOL[BGL-1]-36`, competition ID `2383`) advanced to individual functional and binding assays (visualized on the right bound to the CD20 dimer target as predicted by Chai-1 [20]). **B)** Volcano plot of pooled-screen differential enrichment across all 12,000 competition designs; BAGEL-CAR designs are in teal, with `SOL[BGL-1]-36` labeled in purple. Dashed lines mark the enrichment thresholds: |log_2_ fold change| > 1 (effect size) and false discovery rate (FDR) < 0.05 (multiple-testing-corrected significance). **C–F)** Individual-assay measurements for the top 10 enriched designs across the teams with positive and negative controls. We highlight our `2383` design in color. Asterisks show statistical significance compared to the negative control (***: *p* < 0.001, **: *p* < 0.01). **G)** Surface Plasmon Resonance (SPR) sensorgrams for `2383` binding to CD20.

This paper is organized as follows. First, we lay out our design strategy, showing the three different approaches we have employed, and their varying success rates. Second, we question the refolding validation step and how it would have affected our candidate selection for downstream experimental validation. Finally, we discuss potential reasons for our higher success rate relative to the other approaches tested and suggest how our findings could inform future design campaigns.

## 2 Methods

Our design campaign consisted of three different strategies, schematically presented in Figure 2. We employed BAGEL [18] in all of them, with the recipe scripts provided in the `scripts/car` folder in the `softnanolab/bagel` GitHub repository. To make strategy provenance transparent, we assign each design a BAGEL-CAR identifier in place of the numeric competition IDs.

**Figure 2:**
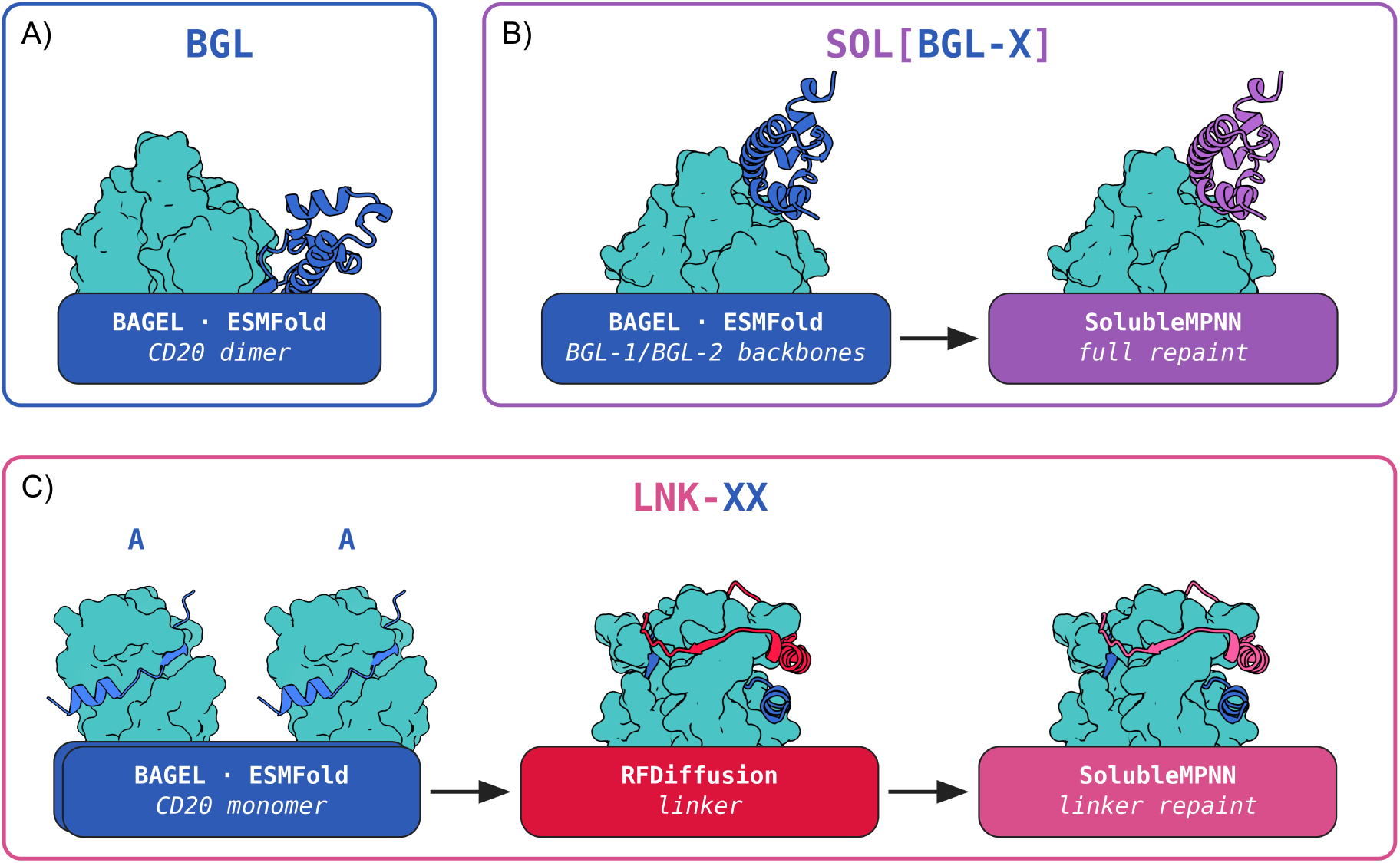
BAGEL-CAR Design Strategies Schematic. **A)** Vanilla 80-residue BAGEL [18] design (`BGL`) against the CD20 dimer (teal). **B)** The two best `BGL` backbones (`BGL-1` and `BGL-2`) fully repainted with SolubleMPNN [21] to increase the number of design submissions while optimizing for solubility and expression. **C)** BAGEL 20-residue binders (`A` and `B`) against individual CD20 monomers, fused via an RFdiffusion-designed linker [10] and repainted with SolubleMPNN to create bivalent constructs (`LNK-XX`).

The first strategy (`BGL`) involves a *vanilla* BAGEL-derived binder against the CD20 dimer. More precisely, we define an energy function in BAGEL that encodes the design objective, and then carry out MC optimization in sequence space to minimize that energy, finding candidate sequences that best satisfy it. This energy is a weighted sum of design terms,

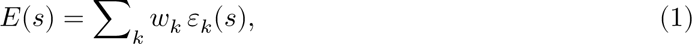

where each *ε_k_* scores a single objective on a candidate sequence *s* and *w_k_* sets its relative weight. To ensure inference tractability, ESMFold [22] is used as the folding oracle that allows us to compute the individual *ε_k_* terms as a function of a predicted structure. More details can be found in Table 1A, and an extensive discussion of each term can be found in Lála et al. [18]. We took the best-performing 50 sequences (ranked by the BAGEL energy) from each of the two independent MC runs and deduplicated them, leaving *n* = 17 unique `BGL` designs, indexed `BGL-1`–`BGL-17` by ascending BAGEL energy. Given the point-mutation nature of our MC algorithm, this yielded low design diversity, motivating us to increase sequence diversity by repainting the sequence (Figure 7, Section S1). Therefore, the two lowest-energy designs, `BGL-1` and `BGL-2`, served as the seed backbones for the next strategy.

**Table 1:** BAGEL Objectives for the Dimer-Targeting (BGL/SOL[BGL-X]) and Monomer-Targeting (LNK-XX) Designs for CD20 Binders. Simulated tempering was used, with temperatures *T*_low_ = 0.1, *T*_high_ = 1.0, and cycle lengths of *n*_low, steps_ = 200, and *n*_high, steps_ = 200. We ran for *n*_cycles_ = 2000. BAGEL’s energy is evaluated using a weighted linear combination based on the terms described in the table, all using ESMFold [22] as the folding oracle. Each row is a term *ε_k_* with weight *w_k_*, combined as in Equation 1. `CD20A` and `CD20B` denote the two CD20 monomers. Epitope and encoding details are given in the Code and Data Availability section.

| A) Dimer Objective (BGL/SOL[BGL-X]) |  |  | B) CD20 Monomer Objective (LNK-XX) |  |  |
| --- | --- | --- | --- | --- | --- |
| *CD20 dimer, 80-residue binder* |  |  | *CD20 monomer, 20-residue binder* |  |  |
| Term | Group | $w_k$ | Term | Group | $w_k$ |
| PTMEnergy | $G_{\text{target}} \cup G_{\text{binder}}$ | 0.2 | PTMEnergy | $G_{\text{target}} \cup G_{\text{binder}}$ | 0.2 |
| PLDDTEnergy | $G_{\text{CD20A}}$ | 1.0 | PLDDTEnergy | $G_{\text{non-epitope}}$ | 0.2 |
| PLDDTEnergy | $G_{\text{CD20B}}$ | 1.0 | PLDDTEnergy | $G_{\text{epitope}}$ | 1.0 |
| PLDDTEnergy | $G_{\text{binder}}$ | 1.0 | PLDDTEnergy | $G_{\text{binder}}$ | 1.0 |
| PAEEnergy | $G_{\text{CD20A}} \leftrightarrow G_{\text{CD20B}}$ | 2.0 | PAEEnergy | $G_{\text{epitope}} \leftrightarrow G_{\text{binder}}$ | 2.0 |
| PAEEnergy | $G_{\text{epitope,A}} \leftrightarrow G_{\text{binder}}$ | 2.0 | SeparationEnergy | $G_{\text{epitope}} \leftrightarrow G_{\text{binder}}$ | 1.0 |
| PAEEnergy | $G_{\text{epitope,B}} \leftrightarrow G_{\text{binder}}$ | 2.0 | HydrophobicEnergy | $G_{\text{binder}}$ | 1.0 |
| SeparationEnergy | $G_{\text{epitopes}} \leftrightarrow G_{\text{binder}}$ | 1.0 | | | |
| HydrophobicEnergy | $G_{\text{binder}}$ | 1.0 | | | |

The second strategy (`SOL[BGL-X]`) repainted these two backbones with SolubleMPNN [21], a solubility-oriented derivative of ProteinMPNN [15]. We label each repainted sequence by its backbone conditioning, for instance, `SOL[BGL-1]-36` derives from `BGL-1`. In addition to improving sequence diversity, repainting biases toward soluble sequences, which the simple sequence-based `HydrophobicEnergy` used above may not optimize sufficiently. The submitted `SOL` set contained 289 `SOL[BGL-1]` and 9 `SOL[BGL-2]` designs, ranked within each group by SolubleMPNN sequence confidence: exp(−ℒ), where ℒ is the mean per-residue cross-entropy of the generated sequence [21].

Third, we tried a more experimental approach (`LNK-XX`) aimed at exploiting multivalent effects by targeting two sites on the CD20 dimer. Instead of designing binders directly against dimeric CD20, we first used BAGEL to generate 20-residue binders against monomeric CD20 using the energy function in Table 1B. From this protocol, we selected the two lowest-energy designs, thereby obtaining two monomer-targeting binders, denoted `A` and `B`, which we combined into three dimer pairings: `AA`, `AB`, and `BB`. We then used RFdiffusion [10] to generate 10 linker backbones per pair and SolubleMPNN to inpaint each backbone with 50 linker sequences. In other words, we redesigned only the linker and kept the monomer-targeting binders fixed to their BAGEL-optimized sequences. To build the input conformation, we superimposed the predicted CD20 monomer from each binder-target complex onto the corresponding chain of the cryo-EM-resolved `6Y97` dimer. The experimental structure thus set the interchain geometry, while the ESMFold-predicted CD20 structure together with the monomer binders was used as input to RFdiffusion and SolubleMPNN. Superimposing the two binder-monomer complexes onto the CD20 dimer places the binder backbones in opposing orientations. In the resulting 80-residue `LNK` fusion, the C-terminal 20-residue segment therefore has the reverse amino acid sequence of its original `A`/`B` binder sequence. For example, `LNK-AB` is `A`-`linker`-`reverse(B)`. As before, we ranked the linker designs by SolubleMPNN sequence confidence, yielding 99 unique linker designs, with a mix of `LNK-AA` (*n* = 14), `LNK-AB` (*n* = 52), and `LNK-BB` (*n* = 33) combinations based on the binder used.

We broadly refer to the strategies as BGL, SOL, and LNK throughout this paper, while sometimes specifying SOL[BGL-X] and LNK-XX to indicate specific sub-strategies.

## 3 Results

### 3.1 Pooled-screen enrichment differed by design strategy

AŁer synthesizing the DNA sequences, primary T cells were engineered to express the CAR library and co-cultured either with CD20-positive Raji cancer cells or without target cells. A design is *recovered* if its barcode was sequenced above the 25-read threshold in all six assay replicates (three CD20+, three no-target controls). A design is *enriched* if it additionally shows statistically significant differential expansion in CD20+ co-culture versus no-target controls, suggesting target engagement by the binder. This enrichment carries two caveats. First, the no-target control establishes only that proliferation depends on the presence of cancer cells, not necessarily that it is CD20-dependent; a CD20-negative target-cell arm would be needed to show the latter. Second, pooled-proliferation enrichment did not cleanly correlate with the individual validation assays in the original data [6], suggesting that second-order effects in pooling can affect binding and functional readouts. Nevertheless, a few designs were fully characterized in the competition. The only one selected from our designs, `SOL[BGL-1]-36`, showed both CD20-specific lysis and CD20 binding by SPR (*K_D_* ≈ 643 nM; Figure 1). With these nuances, we treat enrichment as a proxy for function and compare across teams and design strategies.

As discussed above and shown in Figure 1, BAGEL-CAR designs had the highest enrichment rates in the pooled proliferation screen. In Figure 3, we show that this outcome was strategy-dependent. All strategies containing SolubleMPNN yielded enriched designs in the pooled screen, with hit rates of 19%–56% (`LNK-AB` and `SOL[BGL-2]`, respectively). In contrast, none of the 17 raw `BGL` designs was enriched, although each was recovered by next-generation sequencing (more than 25 reads in all six replicates). Note that recovery measures barcode representation aŁer the screen, not CAR expression. `LNK-AA` and `SOL[BGL-2]` had the highest rates of enriched-if-recovered, with `LNK-AA` having all recovered designs enriched.

**Figure 3:**
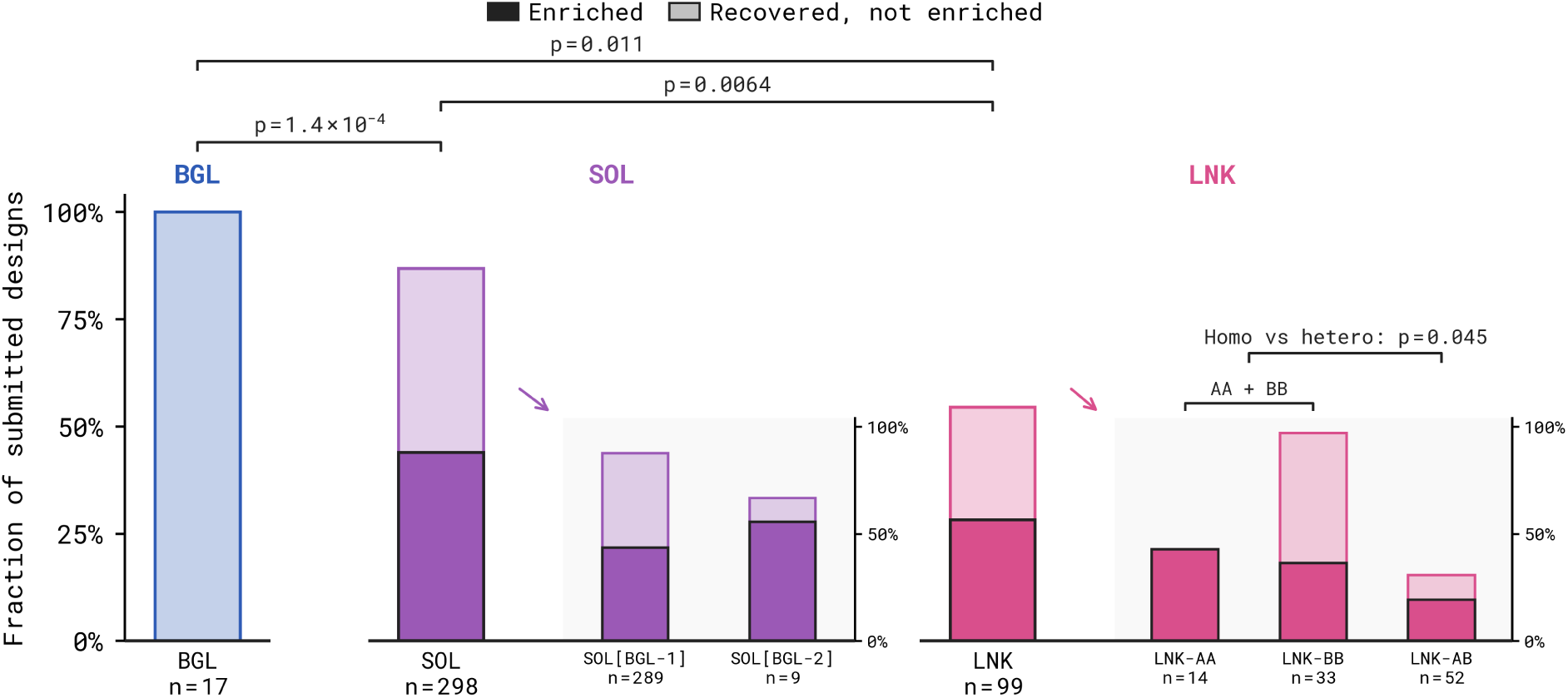
Pooled Proliferation Outcomes by BAGEL-CAR Strategy. Full-height bars show the aggregate `BGL`, `SOL`, and `LNK` strategy outcomes; arrows link `SOL` and `LNK` to half-height axes showing their sub-strategies. Bar height gives recovered / submitted and the solid segment gives enriched / submitted; *n* is the number of submitted designs. Brackets report two-sided Fisher’s exact tests on enrichment (not recovery). The `LNK` comparison pools homo-bivalent `LNK-AA`/`LNK-BB` against hetero-bivalent `LNK-AB`. One design, `SOL[BGL-1]-132`, was significantly depleted and is counted within the recovered-but-not-enriched segment.

SolubleMPNN repainting thus appears necessary for pooled-proliferation enrichment: no raw `BGL` design was enriched (0%), whereas the repainted `SOL` designs were enriched (44%), as were the repainted `LNK` linker fusions (28%), although less oŁen than the `SOL` binders (Figure 3). Within `LNK`, homobivalent (`LNK-AA`/`LNK-BB`) versus heterobivalent (`LNK-AB`) architecture was specifiable *a priori*, whereas we had no prior basis for distinguishing `LNK-AA` from `LNK-BB`. We therefore treat `LNK-AA` and `LNK-BB` as implementations of the same homobivalent strategy; pooled together, they were enriched more oŁen than the heterobivalent `LNK-AB` designs (38% versus 19%).

Two takeaways stand out. First, only designs that used SolubleMPNN – even if only to repaint the linker – were enriched in the pooled assay, while no pure `BGL` design was. Second, the `LNK` fusions showed significant enrichment even though SolubleMPNN repainted only their linker, leaving the binding interface between the monomer-targeting `A`/`B` binders and CD20 carried over directly from ESMFold. Whether this enrichment stems from the ESMFold-derived interface alone, the repainted linker, or both is something we examine in Section 3.3. Unfortunately, given the constraints of the competition, we do not have any data that would measure the function of `A`/`B` binders individually.

### 3.2 ESMFold refolding would have rejected successful designs

As previously discussed, an important self-consistency check for an AI-generated binder is whether the refolded structure engages the same hotspot used as input for the design. In addition to confidence metrics such as ipTM or iPAE, binder-design pipelines therefore also evaluate structural deviation, including binder RMSD and target-aligned pose or interface RMSD [8], [9], [13]. Authors of these methods also sometimes recommend manual inspection to verify that binders remain near the targeted hotspot during generation and refolding. Serendipitously, and only due to competition time constraints, we completely omitted refolding validation aŁer repainting with SolubleMPNN. Therefore, if we had applied this validation step, we would not only have ranked and filtered candidates by model confidence, but also removed designs that did not refold onto the intended epitope, or at least close to that neighborhood. For our submission, this means that we did not verify whether the repainted `SOL` designs or the inpainted `LNK` fusions remained within the intended CD20 hotspot region. Notably, we were motivated to investigate this further aŁer Kosonocky et al. [6] showed that many of the 12,000 designs failed to produce consistent binder placement at the extracellular CD20 epitope. Instead these binders folded against the intracellular side of the membrane protein. Therefore, a natural question occurred to us: if we had included ESMFold, or any other folding algorithm, as a refolding validation step, would we have submitted any of our successful designs?

We approach this question with two complementary geometric measures for each refold. First, as a self-consistency check, we align the predicted CD20 dimer to itself in the input backbone and calculate the binder Cα RMSD relative to the binder in that input backbone (Figure 4A). Large values indicate that the binder has substantially changed its fold or that the prediction places the binder far from its designed pose. Second, to track its global displacement relative to the membrane, we project the center of mass of the binder’s Cα atoms onto the CD20 dimer’s twofold symmetry axis and report its signed membrane-normal coordinate *z*. Throughout this paper, *z* = 0 at the outer (extracellular) membrane leaflet; *z* > 0 lies outside the membrane on the extracellular side, and *z* < 0 extends into and across the membrane. We define the membrane boundaries using Orientations of Proteins in Membranes (OPM) [23]. We do not track the RMSD of the CD20 target dimer here. Although dual orientations exist for some membrane proteins [24], [25], CD20′s orientation seems to be well established, with the extracellular loop 2 (ECL2) epitope facing the extracellular space [26], [27], [28], [29], consistent with the OPM orientation used here.

**Figure 4:**
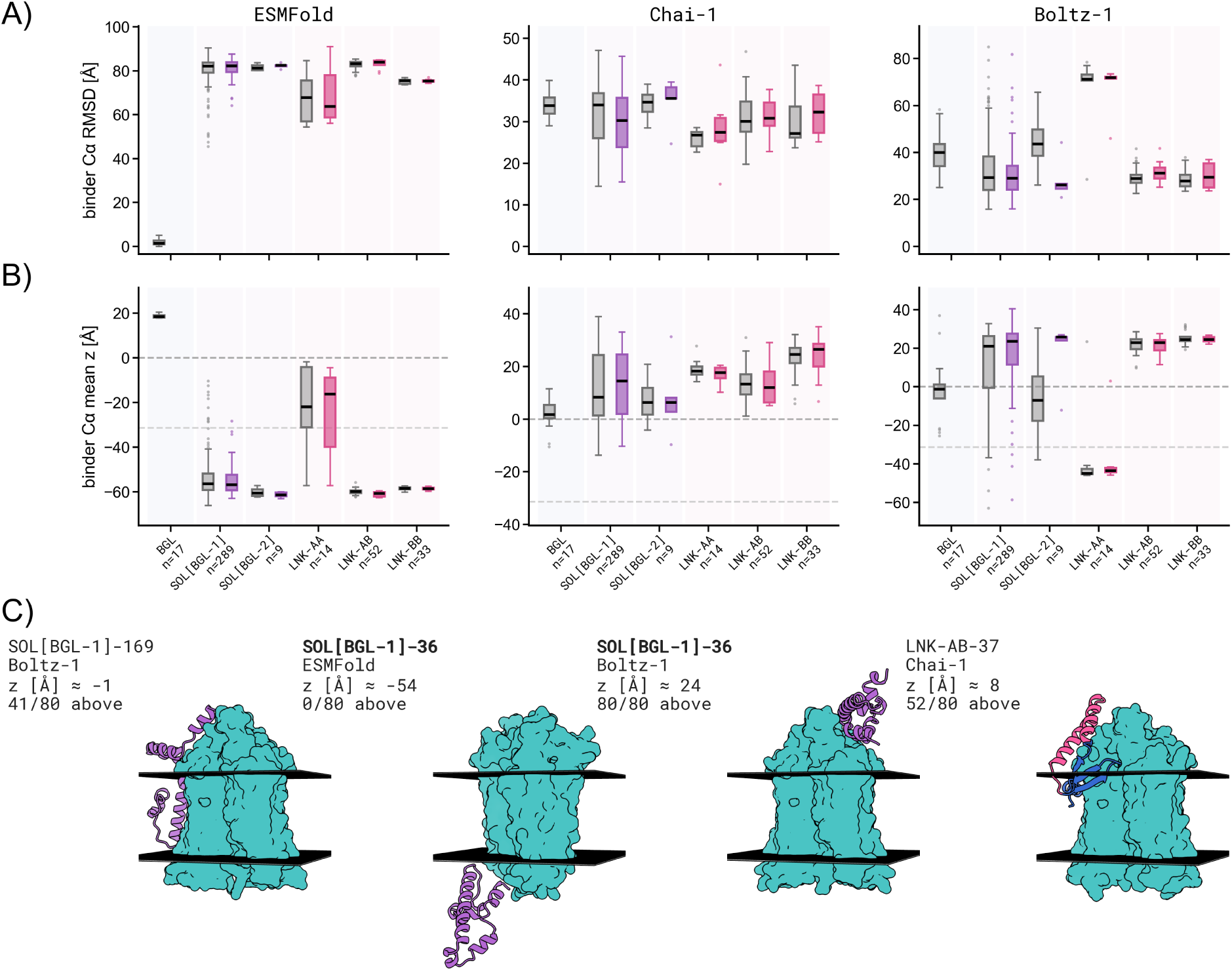
Refolding Analysis of BAGEL-CAR Designs. **A)** Per-strategy self-consistency binder Cα RMSD between the input design backbone (before any repainting if that was part of the strategy) and the refolded prediction. For each model, we align the predicted CD20 dimer directly onto the input design’s own CD20 dimer and compute the RMSD over binder Cα only, thereby measuring RMSD within the target’s frame of reference. **B)** Per-strategy binder Cα center-of-mass *z* along the membrane normal. The horizontal dashed lines mark the outer leaflet at *z* = 0 and the inner leaflet at negative *z*. Boxplots split designs within each strategy by pooled-proliferation outcome, with enriched designs colored and the remaining designs gray; outliers are shown as individual points. Both boxplot rows share the same models and strategies. ESMFold used the BAGEL chain order (`CD20A:CD20B:binder`) with `position_ids_skip=1024` and no explicit glycine linker; Chai-1 used ESM-2 embeddings and no restraints; Boltz-1 used multiple sequence alignments (MSAs). **C)** Representative enriched binder-target complex predictions aŁer alignment to the OPM-oriented CD20 dimer. Binders are colored by design strategy; black planes mark the OPM membrane boundaries. Each render is labeled with the design ID, structure predictor, binder Cα center-of-mass *z* from B), and the number of binder Cα atoms at *z* > 0 out of 80 binder residues. For the validated `SOL[BGL-1]-36` (`2383`), the matched ESMFold and Boltz-1 predictions illustrate their contrasting intracellular and extracellular placements.

Figure 4 summarizes the analysis across ESMFold, Chai-1, and Boltz-1 [30]. First, we see varying performance across the different strategies and refolding oracles. Under ESMFold, `BGL` was the only strategy with consistently extracellular binder placement; predictions for `SOL` and `LNK` placed binders predominantly within or below the membrane. Consistent ESMFold placement is unsurprising since, by construction, `BGL` designs were sampled directly from ESMFold via BAGEL’s MC optimization. On the other hand, although displaced from their original positions,

Chai-1 and Boltz-1 consistently place the binders at the extracellular surface, with some residues still buried in the membrane. `SOL[BGL-1]-169` folded by Boltz-1 and `LNK-AB-37` folded by Chai-1 show examples of such partially buried interfaces, both of which were nonetheless enriched in the pooled screen. Further representative renders are provided in Figure 18 (Section S10). We do not treat visual separation between the enriched and remaining designs in Figure 4 as evidence of outcome discrimination, given the small and unbalanced within-strategy groups.

These results answer the question posed above. Had we applied an OPM-aligned geometric placement filter based on the designed and refolded structures, retaining a design only if more than half of its 80 binder Cα atoms lay at *z* > 0 aŁer CD20 alignment, ESMFold would have rejected all 159 enriched designs, while Chai-1 and Boltz-1 would have rejected 13% and 16%, respectively. This deliberately lenient counterfactual does not apply the model-confidence thresholds usually used to distinguish genuine binding interactions. Kosonocky et al. [6] showed that these metrics poorly discriminated functional designs (see also Figure 17 and Section S9 for a within-team metric screen). We did not test the prospective counterfactual of continuing to generate and refold designs until the filter yielded a full submission set. Given the fraction of candidates that would be removed, especially under ESMFold, this would have required substantially more computational time and resources.

Although the membrane was not represented explicitly during design or refolding, several designs predicted to be partially buried in the membrane were enriched in the pooled screen. We therefore considered whether a hydrophobic binder surface could penetrate the membrane and contact the hydrophobic transmembrane region of CD20, potentially alongside a specific interaction with an exposed epitope. We treat this predicted mechanism cautiously because the predictors disagree on binder placement and their metrics do not provide consistent strategy-local outcome discrimination. Moreover, because the pooled assay lacked a CD20-negative Raji-cell control arm [6], enrichment need not reflect CD20-dependent CAR-T-cell activation. Nonspecific binding to another Raji-cell surface receptor might thus have contributed to the pooled-screen enrichment.

Therefore, we investigate whether exposed binder hydrophobicity supports either possibility in Section S4. Specifically, we fold each binder alone with Chai-1 and define its hydrophobic solvent-accessible surface area (SASA) fraction as the fraction of its exposed surface contributed by apolar residues. We show that higher hydrophobic SASA is associated with Chai-1 membrane-side placement, but the metric does not distinguish designs by either recovery or enrichment status. This provides no evidence that exposed binder hydrophobicity explains pooled-screen proliferation, but it does not exclude specific CD20 transmembrane contacts or nonspecific interactions. Resolving the binding mode experimentally remains outside this study.

One component of the `LNK` designs is that the C-terminal binder segment is reversed in sequence relative to the N-terminal copy. This might affect binding interactions, which we analyze with ESMFold in Section S3 and Figure 9, where the forward and reversed sequences fold at different hotspots. Neither orientation recovers the original extracellular pose. In addition, none of these folding models is chain-permutation invariant, with ESMFold potentially the most sensitive because it lacks native support for multimers. Therefore, for ESMFold, we test whether the conclusion of inconsistent binder placement away from the hotspots is robust to different chain orders and linker encodings in Section S2.

Finally, despite the small sample size (*n* = 14), `LNK-AA` has a perfect enriched-among-recovered rate (100%) and also behaves differently from `LNK-AB` and `LNK-BB` in the ESMFold and Boltz-1 predictions. Under ESMFold, it is the only strategy that places the binder inside the membrane; under Boltz-1, it is the only strategy with an intracellular prediction. However, small-*n* correlations between predicted structure and assay outcome are difficult to interpret causally. The consistent finding is that models disagree and that different design strategies, including the rationally motivated `LNK` linker fusions, can all lead to campaign success. Using ESMFold would have removed all of our enriched submissions, but potentially using some of the other folding models, such as Boltz-1 shown in Figure 4 with our best binder design folding extracellularly, might have worked better. Nevertheless, even if considering Boltz-1 to be the most appropriate refolding predictor, some hits would be still removed (e.g., all `LNK-AA` designs).

### 3.3 Predicted contacts do not identify the source of LNK fusion enrichment

Because repainting `BGL` backbones was associated with enrichment among `SOL` designs, and `LNK` designs were also enriched, we ask whether the diffused-and-repainted `LNK` linker, rather than the ESMFold-derived binding interfaces of the monomer-targeting `A` and `B` binders, was the principal contributor. Although forming an interface was not a linker design objective, it may have been implicitly selected for. As shown in Figure 2C, the diffused linker wraps around the dimer, placing substantial backbone surface in contact with the target.

In Figure 5, we count the contacts of the binder and linker segments of `LNK` to the CD20 extracellular face (not only the narrow hotspot epitope used for BAGEL optimization), before and aŁer the repainting step with SolubleMPNN. For context, the `LNK` designs used 30 different linker backbones (10 per monomer-targeting binder pair), but only six of those survived the final SolubleMPNN score selection. On average, all three sub-strategies lose many of their extracellular contacts relative to their pre-painting input. On average, across both binding segments and the linker, the backbone input contained 38.0 ± 7.8 extracellular contacts; aŁer repainting and refolding, this fell to 0.01 ± 0.1 with ESMFold, 16.1 ± 6.2 with Chai-1, and 12.6 ± 6.3 with Boltz-1. ESMFold predicts no binder contacts with the CD20 extracellular face, whereas Chai-1 and Boltz-1 retain a larger fraction of the initially designed contacts. Under these latter two folding models, the linker itself forms contacts with the target, some of which were already present before repainting. Nevertheless, we see no clear relationship between the number of linker–CD20 contacts and enrichment.

**Figure 5:**
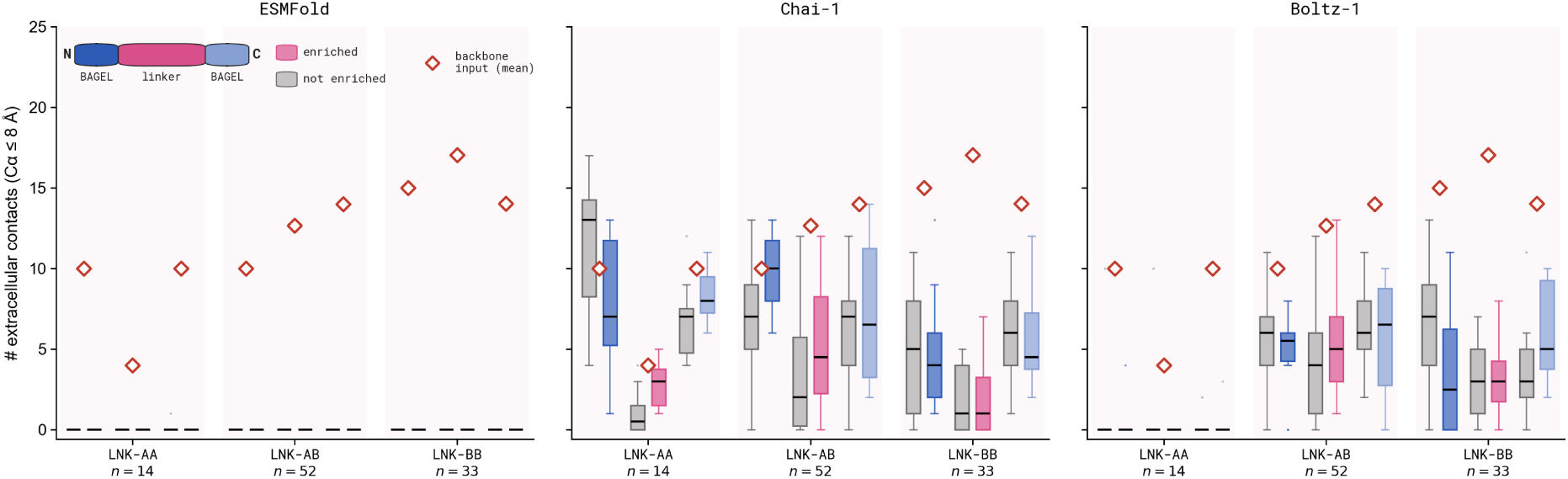
Expected and Predicted Extracellular Contacts for LNK. For each refolded prediction we count binder Cα within 8 Å of any extracellular CD20 Cα (*z* > 0 in the membrane-normal convention of Figure 4; the ECL1 and ECL2 loops that project beyond the membrane), split into the N-terminal BAGEL end (blue), the RFdiffusion linker (pink), and the C-terminal BAGEL end (lighter blue). Red diamonds mark the mean contact count per strategy of the RFdiffusion-diffused backbones (the input to SolubleMPNN). The inset on the top-leŁ of the ESMFold panel shows the coloring scheme for the different parts of the `LNK` designs. Box plots show the per-strategy refolded contacts for the three folding algorithms: ESMFold uses BAGEL chain order (`CD20A:CD20B:binder`, `position_ids_skip=1024`, no glycine linker), Chai-1 uses ESM-2 embeddings and no restraints, and Boltz-1 uses MSAs. The boxplot line shows the median. Designs are pooled by strategy (`LNK-AA`, `LNK-AB`, `LNK-BB`), with *n* denoting the number of designs per strategy, and within each segment shown as the enriched designs (colored) sitting beside the not-enriched designs (gray).

To isolate the contribution of the linker, we also refold the BAGEL monomer-targeting binders against the CD20 dimer without the linker (Figure 9, Section S3). ESMFold predictions do not place the binders at the CD20 extracellular face. Instead, every binder collapses into the membrane or against the intracellular face of CD20 and makes no contacts with extracellular CD20 residues. This might suggest that the linker contributes to binding, although we treat the result cautiously: multichain complex prediction remains a known failure mode, particularly for ESMFold because it lacks native multimer support, and benchmarks have shown reduced interface accuracy as complex size increases from dimers to trimers [22], [31], [32].

We do not evaluate whether the predicted structures correspond to real bound complexes. SolubleMPNN received the linker backbone in close proximity to the target as input, so it remains unclear whether enrichment among `LNK` designs arises from the 20-residue monomer-targeting binders, the diffused-and-repainted linker, or their combination. In Figure 5, the linker tends to form fewer CD20 contacts than the monomer-targeting binders despite being twice as long, although this pattern is not consistent across all sub-strategies. This qualitative result suggests a larger predicted contribution from the binders than from the linker.

### 3.4 ESMFold changes the CD20 target geometry

One key distinction of the ESMFold structures is already visible in Figure 4C, where, for the `SOL[BGL-1]-36` binder, the CD20 target adopts a visibly different geometry from the matched Boltz-1 prediction (the Chai-1 prediction is shown in Figure 18). We investigate this further in Figure 6, where we show that the ESMFold prediction of the CD20 dimer alone adopts a markedly different conformation. This holds across both the *apo* and *holo* states: ESMFold disagrees with the experimental `6Y97` reference, while Chai-1 and Boltz-1 agree closely with it (see Figure 11, Section S5).

**Figure 6:**
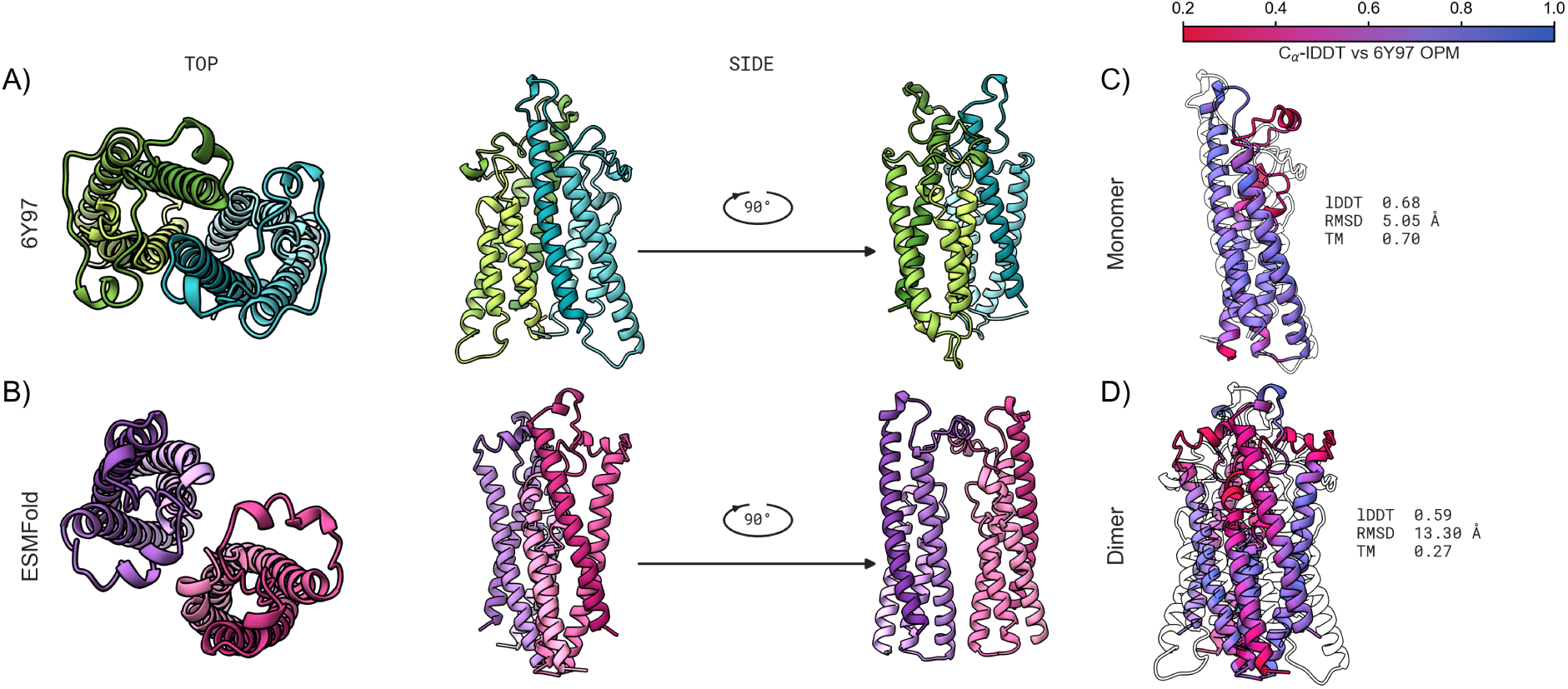
ESMFold Disagrees with the Experimental 6Y97 Structure at Global and Local Scales. **A)** OPM-oriented, cryo-EM-resolved `6Y97` CD20 dimer and **B)** ESMFold target-only prediction (25-glycine linker, `position_ids_skip=512`); three orientations aŁer a single full-dimer Cα superposition – membrane-normal top view (leŁ) and two side views related by a 90° rotation about the membrane normal (right). Chains are colored distinctly, with an N-to-C gradient within each chain. **C)** ESMFold *apo* CD20 monomer and **D)** *apo* CD20 dimer (25-glycine linker), each superposed on the `6Y97`-OPM reference (black silhouette) and colored by per-residue Cα-lDDT [33] (red = poorly preserved, blue = high preservation; 15 Å inclusion radius). In-panel numbers give the global Cα-lDDT, TM-score [34], and Cα RMSD against the reference.

The key difference is a loopy region of one of the transmembrane helices that curls back onto another helix in the `6Y97` structure but is not recovered by ESMFold. This is already visible at the monomer level (Figure 6C), and because all three folding models predict effectively no conformational change upon dimerization – the *apo*-monomer matches the corresponding dimer chain to within ∼1.2 Å Cα RMSD – the same distortion carries directly into the dimer (Figure 6D). This distortion is robust across a sweep over different ESMFold virtual/ glycine linker configurations (Section S8). We believe this difference in the predicted structure of the target to be a dominant aspect, because it affects the binding interface definition. The corresponding per-residue metrics (pLDDT and lDDT) for Chai-1 and Boltz-1 instead confirm that both these models reproduce the experimentally resolved fold almost everywhere (Section S6).

Among all the teams’ methods described in Kosonocky et al. [6], we did not identify another entry that explicitly relied on this ESMFold-predicted CD20 dimer geometry. Given that we used this predicted dimer for all our strategies, and that we achieved high success rates across all but one design strategy, we ask whether this *alternative* conformation contributed to our high hit rate, although we leave this as an open question for future work. Moreover, because no experimentally determined antibody-free CD20 structure is currently available, we cannot distinguish whether ESMFold’s alternative target geometry reflects an unbound CD20 conformer, or a modeling error.

The possibility of an alternative conformation is notable because `SOL` and `LNK` used different repainting approaches (`SOL` was fully repainted, whereas only the linker was repainted in `LNK`) yet both achieved pooled proliferation hit rates of 44% and 28%, respectively. Even if we had removed the more prominent `SOL` designs, the `LNK` strategy would still rank above the 14.8% success rate of the second-best team. This assumes that all other things remain equal in the pooled assay, that there are no second-order effects, and that we do not similarly partition the other teams’ strategies. We also extend this comparison to newer structure predictors in Section S6. In particular, ESMFold2 [35] predicts a CD20 geometry substantially closer to the experimental `6Y97` structure than ESMFold (Figure 13).

Lastly, we want to highlight the nuance of the oligomeric state of CD20 on the cell surface, as that may influence which target geometry is functionally relevant, and should be considered in design campaigns. Resolution enhancement by sequential imaging (RESI) of intact cells shows that untreated CD20 is a mixture of monomers and pre-formed dimers [36]. When antibodies bind, these receptors reorganize into higher-order assemblies whose composition depends on the antibody format. A recent RESI/DNA-PAINT study found that aŁer obinutuzumab (type-II) binding, the population comprised roughly 35% monomers, 48% dimers, 10% trimers, and 7% tetramers, while type-I antibodies such as rituximab and ofatumumab drove more extensive, chain-like oligomerization [37]. For a CAR binder, this means there may not be one obviously “correct” target structure and one oligomeric state: the relevant state is thus whichever oligomer is accessible on the cell surface and still permits CAR clustering, signaling, and formation of a productive immunological synapse [38], [39].

## 4 Conclusion

This retrospective analysis details our approach from the *Bits to Binders* competition. Our validated hit (`SOL[BGL-1]-36`, competition ID `2383`) came from a binder targeting a CD20 dimer, designed by BAGEL and later repainted with SolubleMPNN. Our retrospective analysis identifies SolubleMPNN repainting as associated with pooled-screen enrichment among both `SOL` and `LNK` designs, whereas none of the unrepainted `BGL` designs was enriched. The `LNK` fusions complicate this interpretation: their ESMFold-derived 20-residue binders appeared to retain some BAGEL-optimized contacts, while their diffused-and-repainted linkers formed additional predicted contacts with CD20. In general, the `SOL` (44%) strategy was more successful than `LNK` (28%), but `LNK` was still successful enough to have remained the top-scoring proliferation strategy in the entire competition. We observed that using ESMFold-derived CD20 structures in both monomeric and dimeric complexes, rather than the experimentally resolved `6Y97` structure or the closely matching Chai-1 and Boltz-1 predictions, might have contributed to our high hit rate. The key methodological observation, however, is that the standard backbone-inpainting-validation pipeline would have led us to reject our successful designs. With ESMFold as the refolding oracle, we would have rejected all 159 enriched designs, whereas Chai-1 and Boltz-1 would have rejected 13% and 16%, respectively. As noted, pooled enrichment is an imperfect proxy for individual binding performance, so these conclusions should be interpreted with that caveat in mind, but we find it promising to see BAGEL-centric campaigns perform competitively. We therefore suggest that protein design pipelines should be stress-tested on specific problems, with intermediate outputs from filtering stages also tested rather than relying on *common practices* inherited from the community. Lastly, we see this as only the starting point in a wider effort to apply novel approaches of computational protein design to specific and effective cell therapy treatments.

## Author Contributions

J.L. and S.A.-U. contributed equally to all aspects of this work.

## Code and Data Availability

The original *Bits to Binders* competition data (design sequences, pooled-screen enrichment results, and validation assays) are released by Kosonocky et al. [6] on GitHub at `github.com/ kosonocky/bits-to-binders`. The BAGEL design framework is available on GitHub at github.com/softnanolab/bagel, with the BAGEL-CAR recipe scripts in the `scripts/car` folder. Structure prediction was carried out through `boileroom` [40], a unified interface for protein-structure folding models, available on GitHub at github.com/softnanolab/boileroom. The released scripts use `position_ids_skip=512` with no glycine linker, a minor deviation from the original campaign value of `1000` that we do not expect to affect the qualitative interpretation. The BAGEL target sequence was the 169-residue CD20 segment resolved in `6Y97` (UniProt residues 45–213), reindexed to 1–169 for the design input. The monomer-seed objective and `CD20A` in the dimer objective used input residues His114–Asn132 (HTPYINIYNCEPANPSEKN), corresponding to UniProt residues His158–Asn176. `CD20B` used input residues His114–Lys131 (UniProt His158–Lys175), an accidental one-residue asymmetry preserved in the released scripts for faithfulness to the competition run.

## Acknowledgements

We are deeply thankful to all the organizers of the *Bits to Binders* competition, namely Clayton W. Kosonocky, Alex M. Abel, Aaron L. Feller, Amanda E. Cifuentes Rieffer, Phillip R. Woolley, Daryl R. Barth, Tynan Gardner, Stephen C. Ekker, Andrew D. Ellington, Wesley A. Wierson, and Edward M. Marcotte, for designing and running this unprecedented community benchmark. In particular, we thank Clayton W. Kosonocky and Phillip R. Woolley for valuable discussions and assistance with post-competition data analysis. We also thank the competition sponsors and partners: LEAH Laboratories, Adaptyv Bio, Twist Bioscience, the University of Texas at Austin Center for Systems and Synthetic Biology, the Texas Advanced Computing Center (TACC), Modal, Lonza, ScaleReady, KUNGFU.AI, Nucleate AI in Biotech, and The BioML Society. Lastly, we acknowledge our team member David Miller who took part with us in the early stages of the competition.

## Supplementary Information

### S1 Design strategy sequence diversity

We computed pairwise sequence distances within each design strategy using three metrics: Hamming distance (fraction of mismatched positions), BLOSUM62-derived distance [41] (substitution severity), and Grantham distance [42] (physicochemical property difference). All three agree qualitatively, as shown in Figure 7. The 17 `BGL` designs originate from two independent BAGEL MC runs (`BGL-MC1`, *n* = 10; `BGL-MC2`, *n* = 7). Each run converged to a tight sequence neighborhood (within-run median Hamming 0.04–0.23), but the two neighborhoods share almost no sequence identity (93% of positions differ). For `LNK` strategies, the first and last 20 residues are the fixed BAGEL-derived monomer binders, so distances are computed only over the 40-residue RFdiffusion/SolubleMPNN linker region. On this basis, `LNK-AB` is the most diverse strategy (median Hamming 0.73), while `LNK-BB` and `LNK-AA` show moderate linker diversity (median 0.25 and 0.33). `SOL` designs fall in a similar range (median 0.20–0.31).

**Figure 7:**
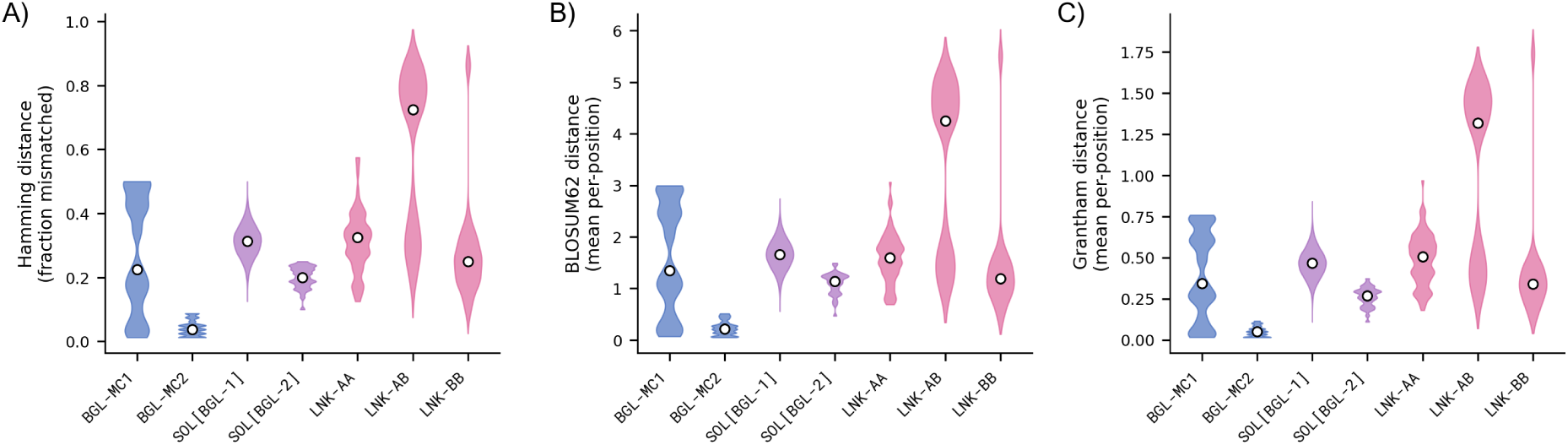
Design Strategy Sequence Diversity. Distribution of pairwise distances among all designs within each strategy. `BGL` designs are split by their two source MC runs (`BGL-MC1`, *n* = 10; `BGL-MC2`, *n* = 7), which converged to unrelated sequence neighborhoods. For `BGL` and `SOL` strategies, distances are computed over all 80 residues; for `LNK` strategies, only over the 40-residue linker region (the flanking 20-residue monomer binders are identical within each strategy). **A)** Hamming distance (fraction of mismatched positions). **B)** BLOSUM62-derived distance [41] (mean per-position substitution cost). **C)** Grantham distance [42] (mean per-position physicochemical distance). White dots mark the median.

### S2 ESMFold chain-order sensitivity

In Figure 8, we evaluate whether the chain order passed to ESMFold affects how many binders fold on the correct side of the target. We compare three placements of the binder within the `CD20A`:`CD20B`:`binder` triplet: `start` (`binder`:`CD20A`:`CD20B`), `mid` (`CD20A`:`binder`:`CD20B`), and `end` (`CD20A`:`CD20B`:`binder`, the BAGEL recipe). We run each of these settings with and without a 25-glycine linker, while always using a positional skip of 1024 indices. “Correct side” is defined as at more than half of the binder Cα atoms lying above the extracellular membrane leaflet of the OPM-aligned reference. We show that only a few of the `BGL` designs would pass, along with some `SOL[BGL-2]` and `LNK-AA` designs. The conclusions from Figure 4 are insensitive to chain order and glycine-linker encoding.

**Figure 8:**
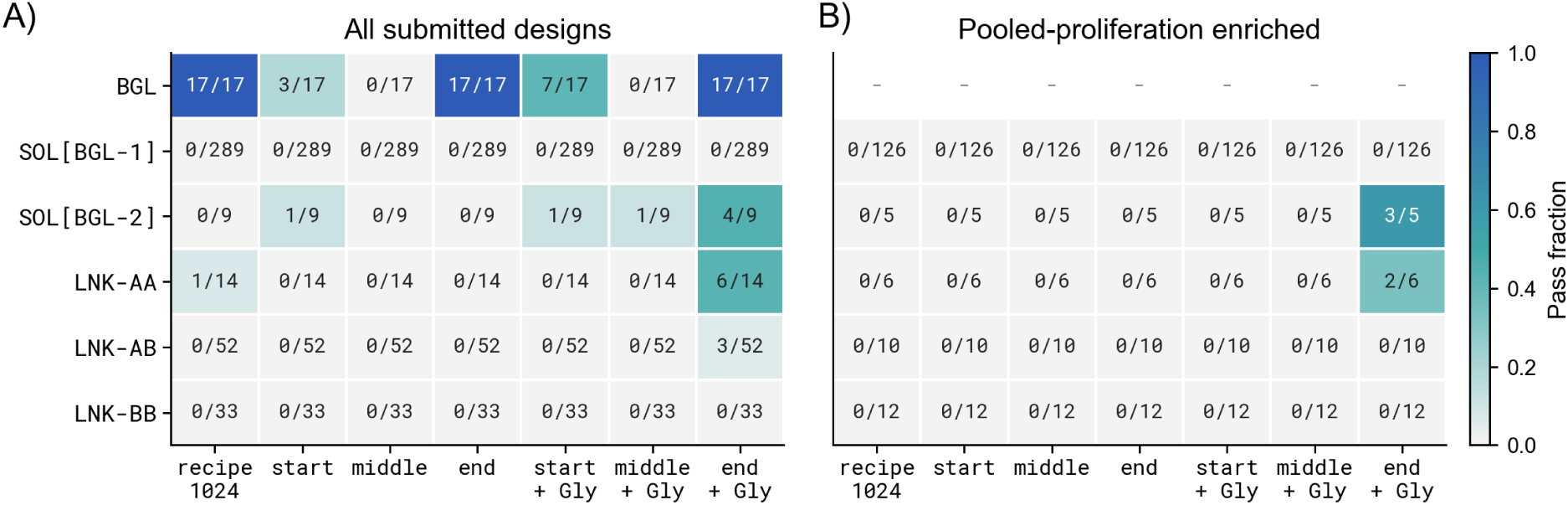
ESMFold Binder Placement Sensitivity to Chain and Linker Encoding. Pass counts under the rule used in Figure 4: more than half of binder Cα atoms at *z* > 0 aŁer CD20 alignment. Columns compare the BAGEL-chain-order setup (`CD20A`:`CD20B`:`binder`, `position_ids_skip=1024`, no glycine linker) with `start` (`binder`:`CD20A`:`CD20B`), `middle` (`CD20A`:`binder`:`CD20B`), and `end` (`CD20A`:`CD20B`:`binder`) encodings, each without or with an explicit glycine linker. The 25-glycine linker is denoted with `Gly`. **A)** All 414 submitted designs. **B)** Pooled-proliferation-enriched designs only; the empty `BGL` row reflects that no unrepainted `BGL` design was enriched.

### S3 Monomer-targeting LNK binders with no linker

**Figure 9:**
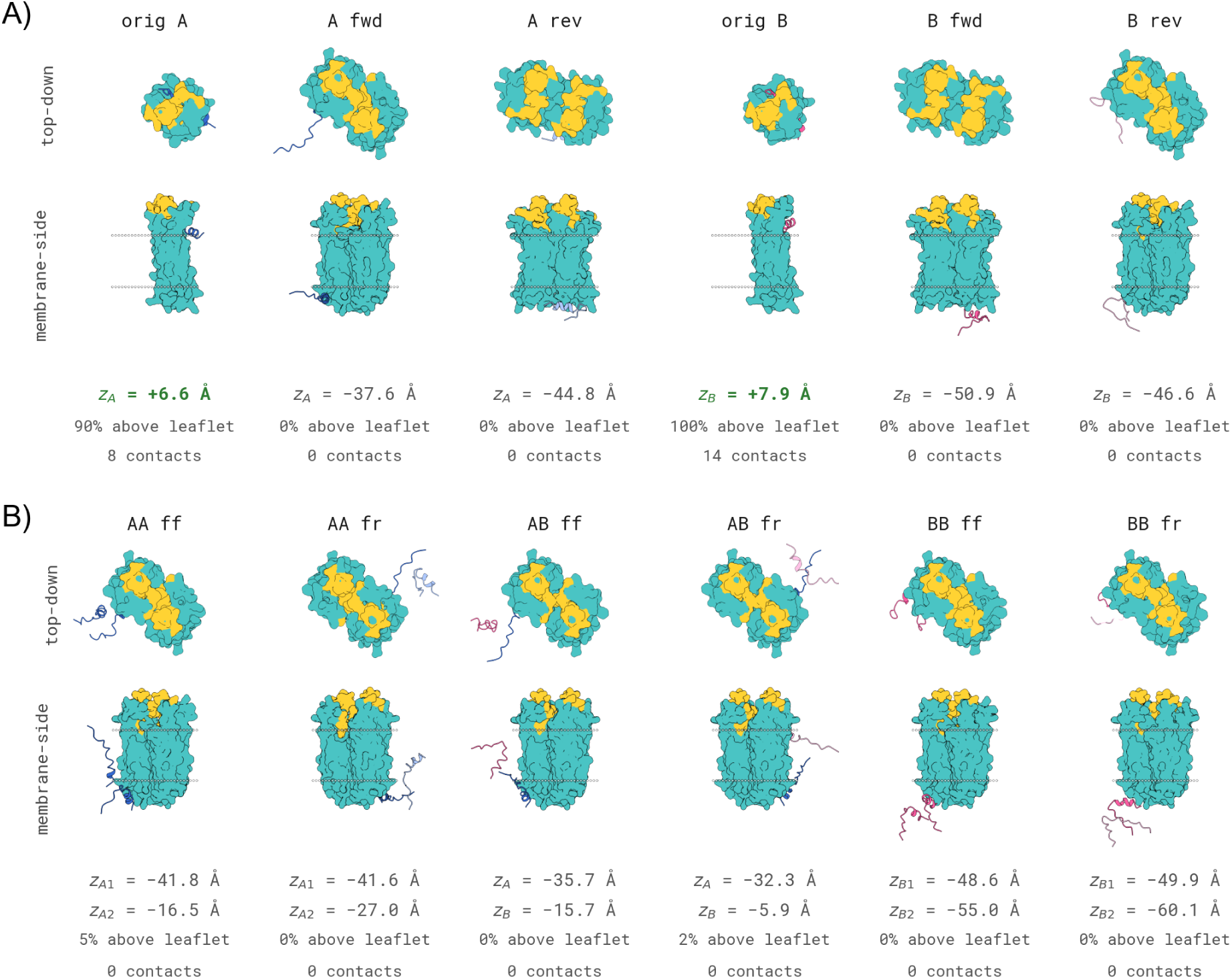
Linkerless LNK Monomer Halves and Their Original Reference Poses. Structures are shown in two membrane-oriented views (top-down extracellular face and membrane-side). The `orig A` and `orig B` columns are the original as-designed BAGEL monomer-complex poses, not ESMFold refolds; all remaining columns are ESMFold predictions against the CD20 dimer. CD20 is shown in teal with the original target epitope in yellow. Binder A is shown in blue, binder B is shown in red, and reversed copies are shown in lighter tints. Gray spheres mark the membrane leaflets (membrane-side view only). Below each construct, *z* is the mean binder Cα membrane-normal coordinate, with *z* = 0 at the outer leaflet and *z* > 0 extracellular; positive values are shown in green and bold. The `A` or `B` subscript identifies the BAGEL-derived 20-residue binder being measured. For `AA` and `BB`, the additional `1` and `2` distinguish the first and second binder chains in the ESMFold input. We also report the percentage of pooled binder Cα at *z* > 0 and the number of pooled binder residues contacting the CD20 extracellular face (binder Cα within 8 Å of any extracellular CD20 Cα, also defined by *z* > 0; contact definition as in Figure 5). **A)** The original `A` and `B` reference poses alongside each isolated 20-residue seed refolded in forward (`f`) and reversed (`r`) sequence orientations. **B)** The two BAGEL-derived binder segments refolded together without a linker.

### S4 Binder hydrophobic SASA and Chai-1 membrane placement

For each Chai-1 binder-alone prediction, we calculated atom-level solvent-accessible surface area using a 1.4 Å solvent probe. We define the hydrophobic SASA fraction as the summed SASA of apolar residues (A, V, I, L, M, F, W, Y, and P) divided by the total binder SASA. Across all 414 designs, this fraction correlated with more membrane-side placement (binder Cα atoms at or below the extracellular leaflet: Spearman *ρ* = 0.501; binder mean z: *ρ* = −0.484; both permutation *p* ≤ 1.0 × 10^−4^). It did not distinguish pooled-screen enrichment (Mann–Whitney *p* = 0.36), and the recovery-based comparisons in Figure 10A and Figure 10B were also not significant aŁer Holm correction (both *p* = 0.065). Because both the binder-alone folds and placement predictions come from Chai-1, this analysis tests a model-internal relationship, not binding, specificity, or mechanism.

**Figure 10:**
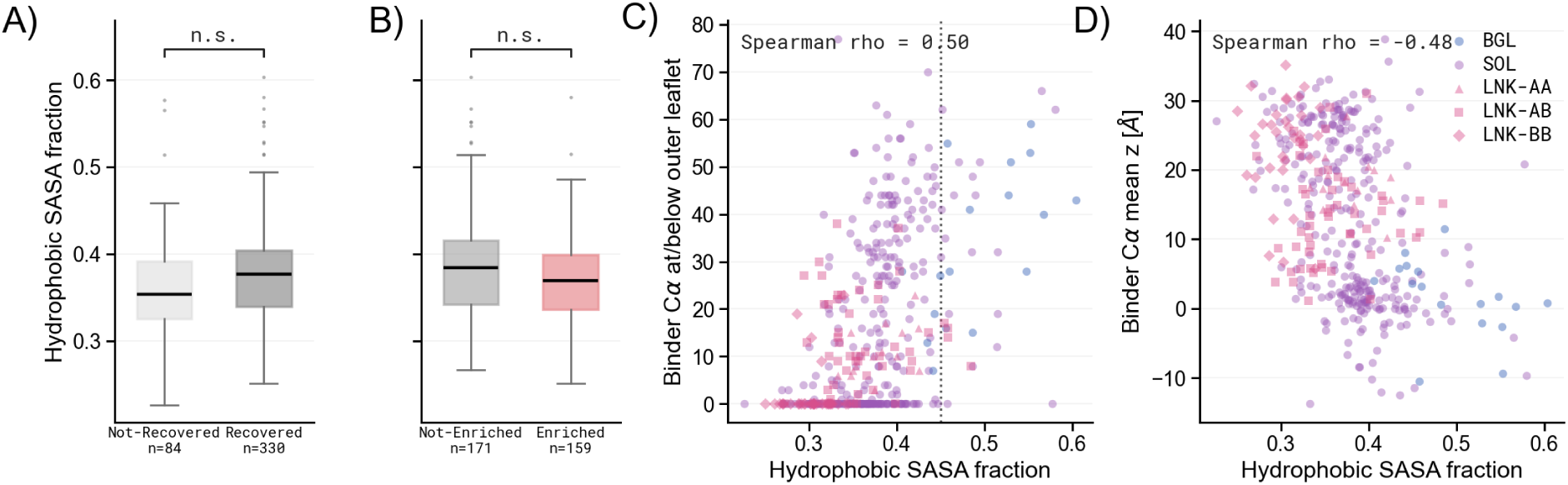
Binder Hydrophobic SASA Across Recovery, Enrichment, and Chai-1 Membrane Placement. **A)** Hydrophobic SASA fraction for 330 recovered and 84 non-recovered designs. **B)** Among recovered designs, hydrophobic SASA fraction for 159 enriched and 171 not-enriched designs. Brackets in **A)** and **B)** report the Holm-corrected status of the two exploratory comparisons. **C)** Hydrophobic-residue SASA fraction from Chai-1 binder-alone structures versus the number of binder Cα atoms at or below the extracellular outer leaflet. The vertical guide at 0.45 captures 145 of the 147 designs with all 80 binder Cα atoms above the outer leaflet, but is not a strict cutoff. **D)** The same metric versus binder Cα mean *z*, using the convention defined in Figure 4. **C)** and **D)** use the Chai-1 `none` OPM-frame placement table for all 414 BAGEL-CAR designs; reported correlation *p* values are at the 10,000-permutation floor.

### S5 Target-dimer geometry across refolding strategies

The qualitative split between ESMFold, Chai-1 and Boltz-1 in Figure 6 holds across the full design portfolio. Figure 11 extends the comparison to the 414 BAGEL-CAR *holo* (i.e., bound in the binder-target complex) predictions, scoring each predicted CD20 dimer against the OPM-aligned experimental `6Y97` reference. ESMFold remains far from the reference across all chain orderings and different linker encodings, while Chai-1 and Boltz-1 stay tight to the *apo*baseline. Lastly, we do not investigate the apparent shiŁ to lower RMSD and higher TM-score when using a glycine linker with the `start` or `end` settings.

**Figure 11:**
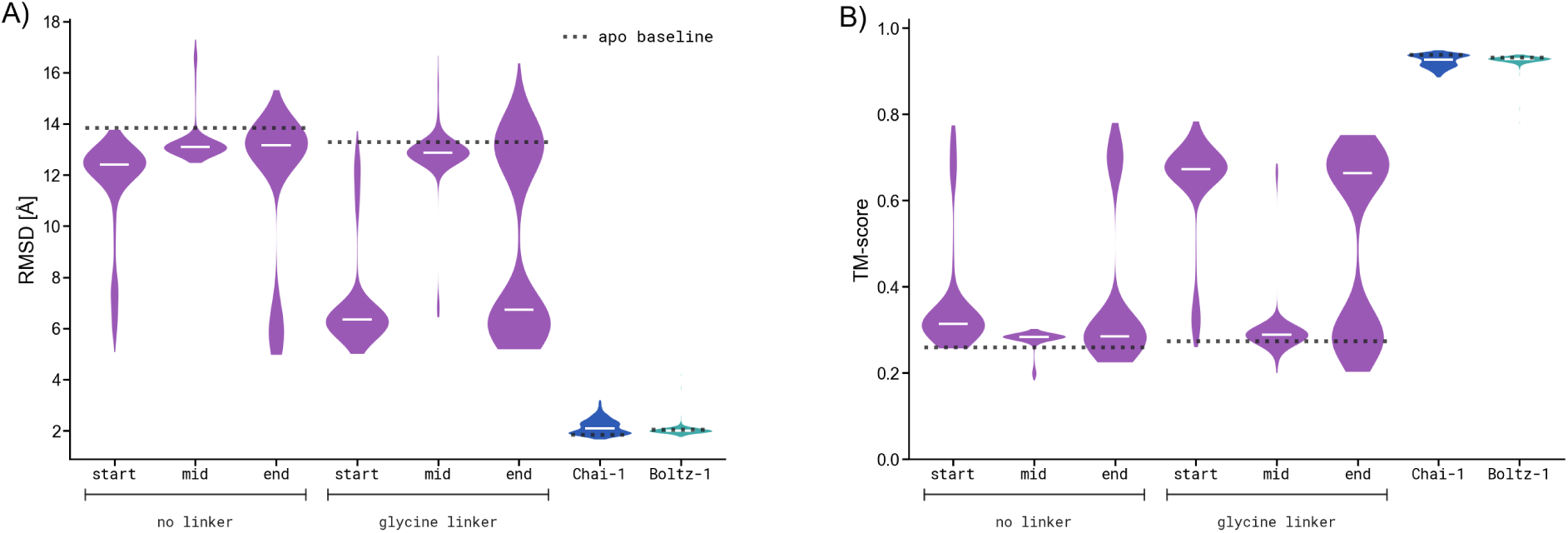
Target-Dimer Geometry of *Holo* CD20 Predictions Across Refolding Strategies. **A)** CD20 target-dimer Cα RMSD (Å) and **B)** TM-score [34] against `6Y97` for the 414 BAGEL-CAR *holo* complex predictions, taking the lower-RMSD of the two symmetric chain mappings. ESMFold predictions are split by linker encoding (no linker vs. 25-glycine linker) and binder chain position within the triplet – `start` (`binder`:`CD20A`:`CD20B`), `mid` (`CD20A`:`binder`:`CD20B`), `end` (`CD20A`:`CD20B`:`binder`); Chai-1 (with ESM-2) and Boltz-1 (with MSA) are shown to the right. Each violin is filled with the predictor color; thin white bars mark per-group medians. Black dotted lines indicate the corresponding target-only *apo* baselines.

### S6 Per-residue C_α_-lDDT of *apo* CD20 predictions

The per-residue C_α_-lDDT decomposition of the ESMFold *apo* predictions, which localizes ESMFold’s disagreement with the OPM-aligned `6Y97` reference to the BAGEL-epitope-bearing extracellular loop, is shown in Figure 6C and Figure 6D. Here we report the corresponding diagnostic for Chai-1 and Boltz-1 (Figure 12). Both reproduce the OPM fold almost everywhere in both monomer and dimer contexts (global C_α_-lDDT 0.77–0.80), with low-lDDT residues confined to the flexible cytoplasmic N-terminal tail.

**Figure 12:**
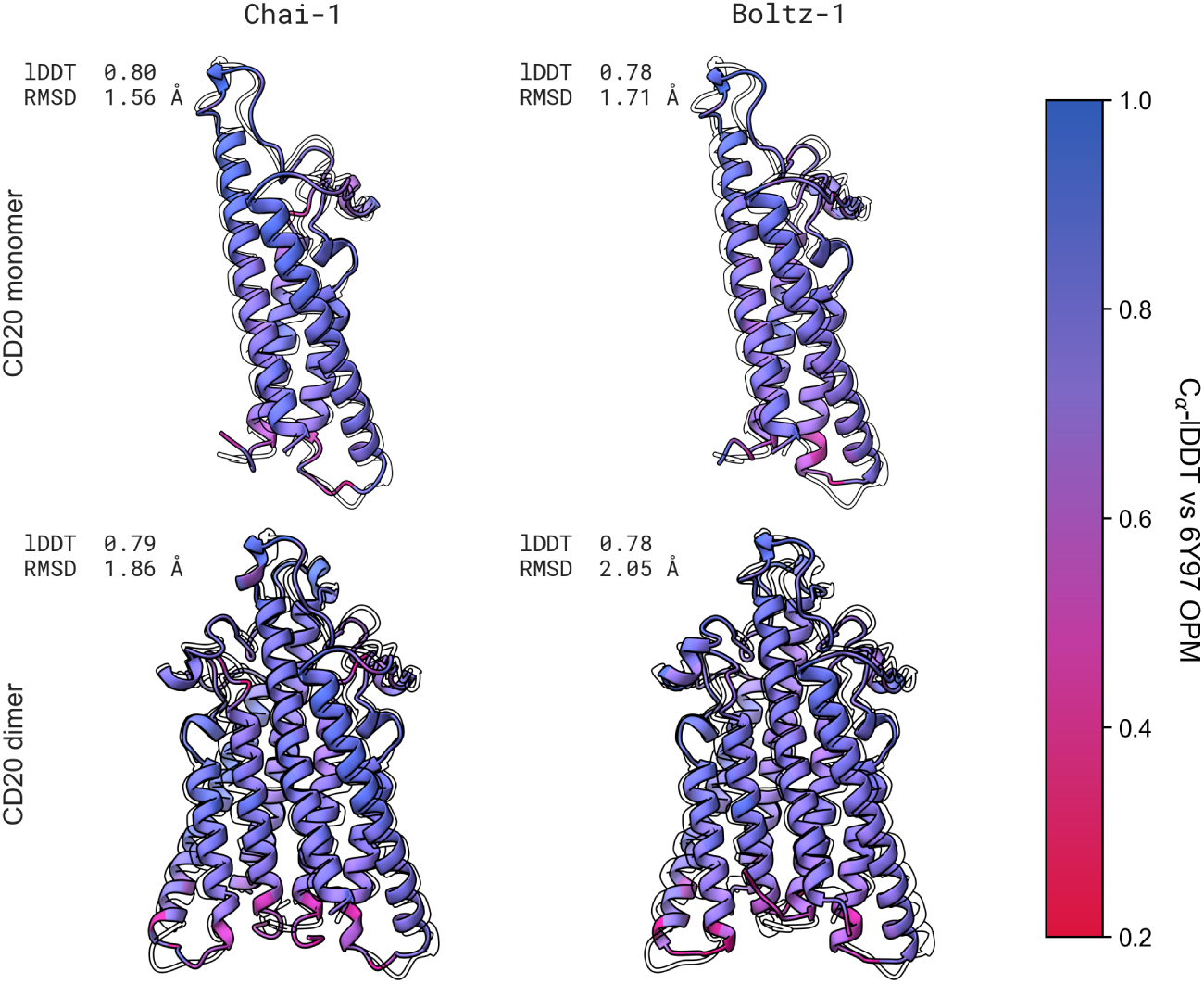
Per-Residue Cα-lDDT of *Apo* CD20 Predictions Against the 6Y97 OPM Reference. Rows are monomer vs. dimer scope; columns are folding models (Chai-1, Boltz-1). Each cartoon is the prediction superposed on the 6Y97 OPM reference (black silhouette) and colored by per-residue Cα-lDDT [33] (red = poorly preserved local environment, blue = high preservation; inclusion radius 15 Å). In-panel numbers report global Cα-lDDT and Cα RMSD. The ESMFold counterparts are shown in Figure 6C and Figure 6D.

For completeness, we also expand the analysis here using newer models - specifically, ESMFold2 [35], Boltz-2 [43], and Protenix [44]. Figure 13 shows that most models recapitulate the experimental `6Y97` structure in both monomeric and dimeric contexts. Protenix and Boltz-2 fail in the displayed no-MSA configurations, whereas ESMFold2′s language-model-based mode does not degrade without an MSA. Most notably, ESMFold2 recovers the experimental `6Y97` structure much more closely than ESMFold. Whether this closer agreement reflects improved structural accuracy or familiarity with experimentally resolved structures represented in its training data, rather than the exclusion of a physically plausible alternative conformation, remains open for further study.

**Figure 13:**
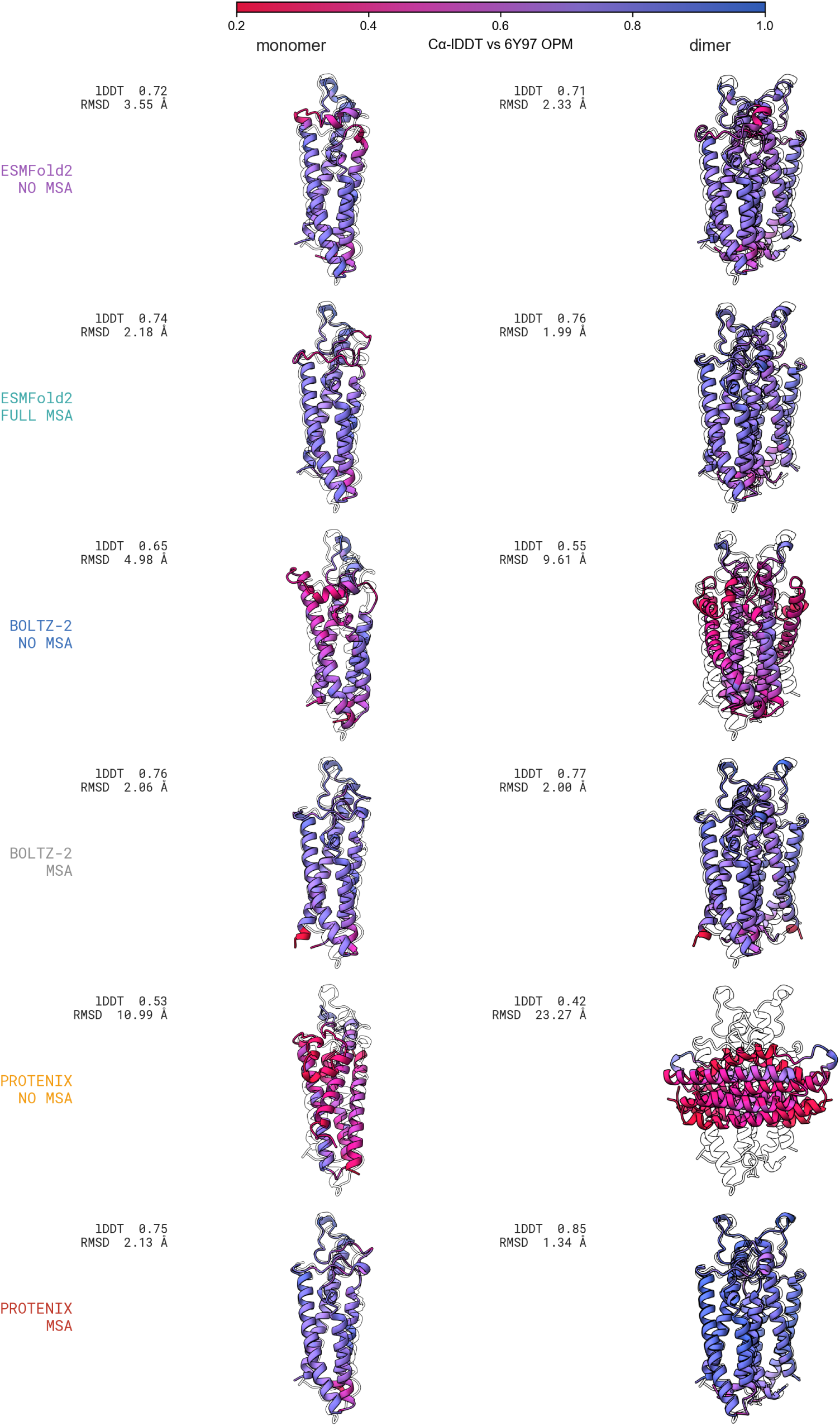
CD20 Geometry Across Newer Structure Predictors. Target-only CD20 monomer (leŁ) and dimer (right) predictions from ESMFold2, Boltz-2, and Protenix superposed on the OPM-aligned `6Y97` reference (gray outline) and colored by per-residue Cα-lDDT. Rows show the evaluated model configurations, with each cell displaying the selected sample. In-panel numbers report global Cα-lDDT and Cα RMSD against `6Y97`.

Figure 14 complements the render above with per-residue traces of each model’s confidence (pLDDT, colored by model) against the ground-truth C_α_-lDDT (black). ESMFold and Chai-1 pLDDT track lDDT across both contexts; Boltz-1 is the exception, with lower pLDDT that under-represents per-residue lDDT, most clearly in the monomer. For ESMFold, this correspondence suggests that the model reflects its own uncertainty in the distorted regions. Whether this represents genuine conformational flexibility or well-calibrated confidence in a poor prediction is unclear from these data alone and would require experimental investigation. We also note that both ESMFold’s confidence (pLDDT) and the actual lDDT are lower in the dimer than in the monomer, potentially arising from the lack of native multimeric support.

**Figure 14:**
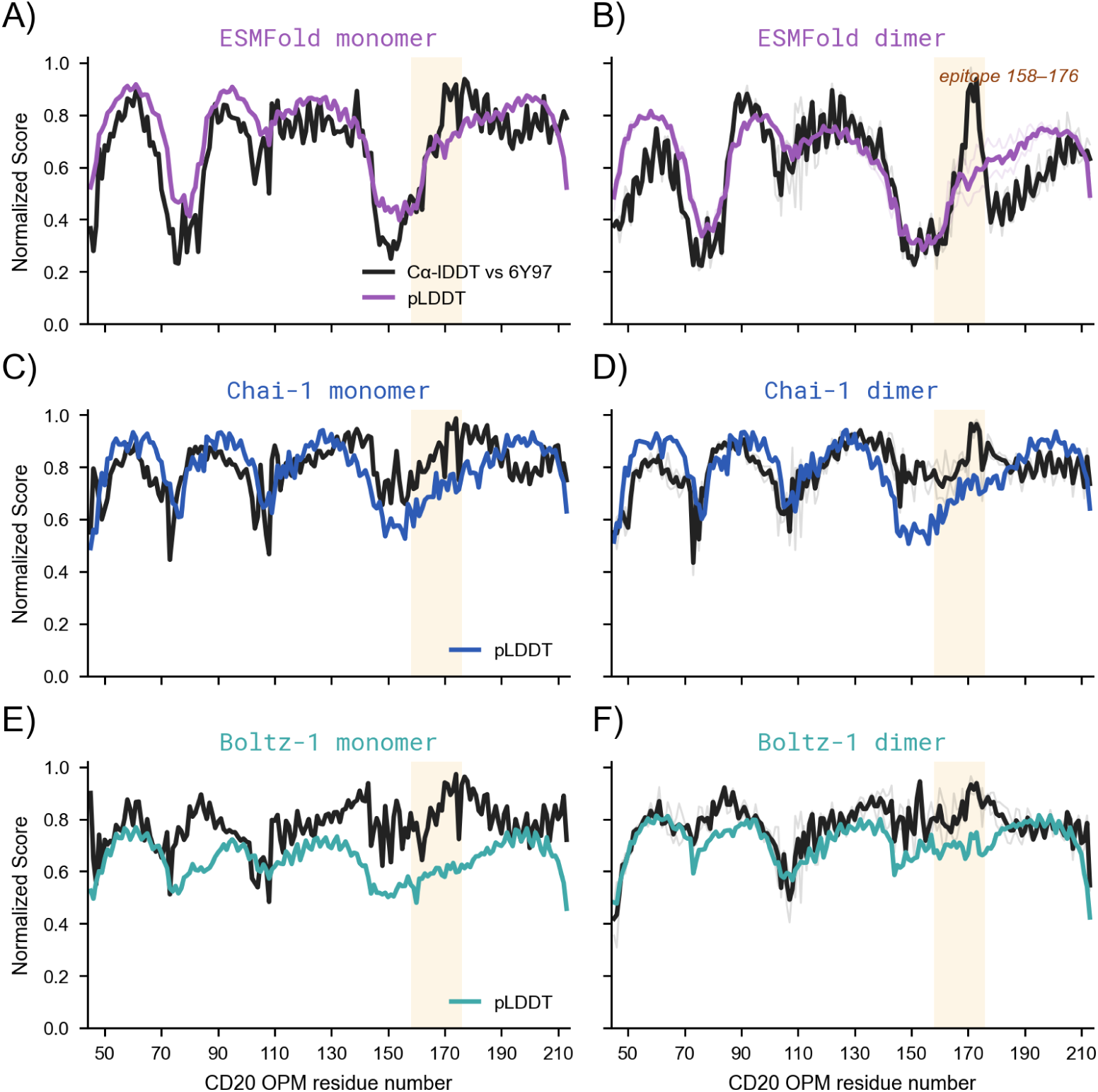
Per-Residue pLDDT and Cα-lDDT of *Apo* CD20 Predictions. Rows are folding models (ESMFold, Chai-1, Boltz-1); columns are monomer vs. dimer scope. Black traces show per-residue Cα-lDDT against the OPM-aligned `6Y97` reference. Colored traces show each model’s pLDDT normalized. Faint traces are individual monomeric chains in the dimer. Thick traces are averaged across both chains. Orange band marks the used epitope in BAGEL (residues 158–176).

### S7 Predicted CD20 monomer-vs-dimer conformational shift

The similar model-specific distortions in the monomer and dimer predictions raise a narrower question: does dimerization itself alter the predicted fold of each CD20 chain? We compare each monomer prediction against the lower-RMSD chain of the corresponding dimer prediction. All three folding models give 1.1–1.2 Å Cα RMSD, below the 1.47 Å chain-A-vs-chain-B asymmetry within the experimental `6Y97` dimer (Table 2). In other words, we do not see a substantial structural shiŁ upon dimerization, and the monomers effectively remain rigid.

**Table 2:**
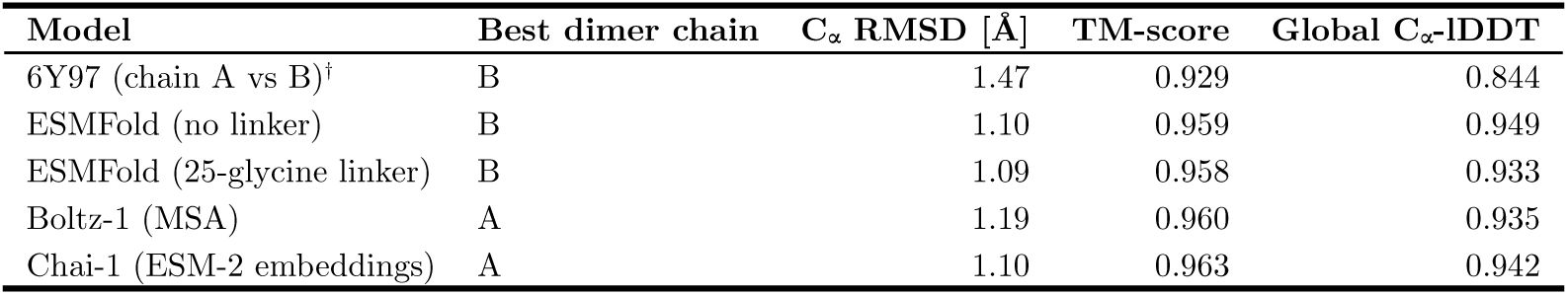
Predicted CD20 Monomer-vs-Dimer Conformational Shift. For each folding model, the *apo*-monomer Cα coordinates are superposed via the Kabsch algorithm onto one chain of the *apo*-dimer prediction over the longest exact-sequence overlap; we report the lower-RMSD chain of chains A and B. TM-score [34] is normalized by the monomer Cα length; global Cα-lDDT uses R0 = 15 Å and thresholds {0.5, 1, 2, 4} Å with the monomer as the reference. ^†^ No experimentally resolved full-length CD20 *apo* monomer is available, so the `6Y97` row reports the intra-dimer asymmetry between chain A and chain B of the OPM-oriented structure as a noise-floor baseline rather than a true monomer-vs-dimer comparison.

### S8 ESMFold chain-encoding sweep for the CD20 dimer

We refold the *apo* CD20 dimer across `position_ids_skip` ∈ {0, 256, 512, 1024, 2048} and glycine-linker length ∈ {0, 5, 25, 50} to check whether the ESMFold-vs-`6Y97` disagreement in Figure 6 is an encoding artifact (Figure 15). 18 of 20 cells cluster at C_α_ RMSD 12.7–13.9 Å (TM-score 0.26–0.30), including the Figure 6 reference (`position_ids_skip=512`, 25-glycine linker) and the BAGEL recipe (`position_ids_skip=1024`, no linker). Only `position_ids_skip=0` with linker ≤ 5 recovers a substantially different geometry (RMSD 7.13–8.44 Å, TM-score 0.62–0.67). Within the dimer regime, the skip value saturates beyond the first nonzero column. Inspection of the `transformers` [45] ESMFold implementation used inside `boileroom` [40] uncovers two compounding causes: `position_ids` is forwarded only to ESMFold’s folding trunk and structure module, never to the ESM-2 encoder – which therefore sees the two CD20 copies as one contiguous sequence regardless of skip value – and the folding trunk’s pairwise relative-position embedding is one-hot binned and clipped at ± 32 residues, a value inherited verbatim from AlphaFold2′s `max_relative_feature` ([46], Algorithm 4). We treat the missing ESM-2 position-ID routing as a general bug. Boileroom version 0.3.1 corrected it for direct ESM-2 inference, although the ESMFold wrapper used here remains outside that fix. By contrast, clipping the pairwise relative-position feature at ± 32 is an inherited ESMFold implementation detail: separations beyond 32 positional indices map to the same terminal bin, so larger linker- or skip-induced offsets are indistinguishable to this part of the folding trunk.

**Figure 15:**
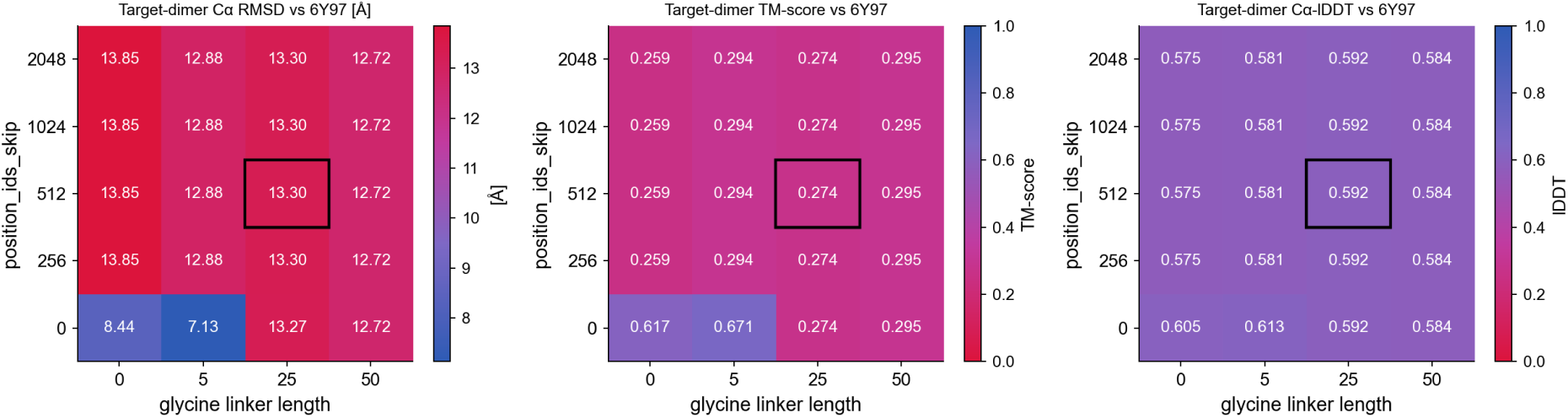
ESMFold *Apo* CD20 Dimer Geometry Across Chain-Encoding Hyperparameters. Target-dimer Cα RMSD (Å, leŁ) and TM-score [34] (right) of ESMFold *apo* CD20 dimer predictions against the OPM-aligned `6Y97` reference, as a function of `position_ids_skip` (rows) and glycine-linker length (columns). The Figure 6 reference cell (`position_ids_skip=512`, 25-glycine linker) is boxed in black. The bottom-leŁ island (`position_ids_skip=0`, linker length ≤ 5) is the only regime that recovers a near-reference fold.

We visualize the two low-RMSD cells and the `position_ids_skip=1024`, no-linker cell using the per-residue Cα-lDDT treatment from Figure 6D (Figure 16). The two low-RMSD predictions track the OPM reference across the membrane core (global Cα-lDDT 0.60–0.61), whereas the `position_ids_skip=1024`, no-linker prediction is displaced within the membrane plane (RMSD 13.85 Å).

**Figure 16:**
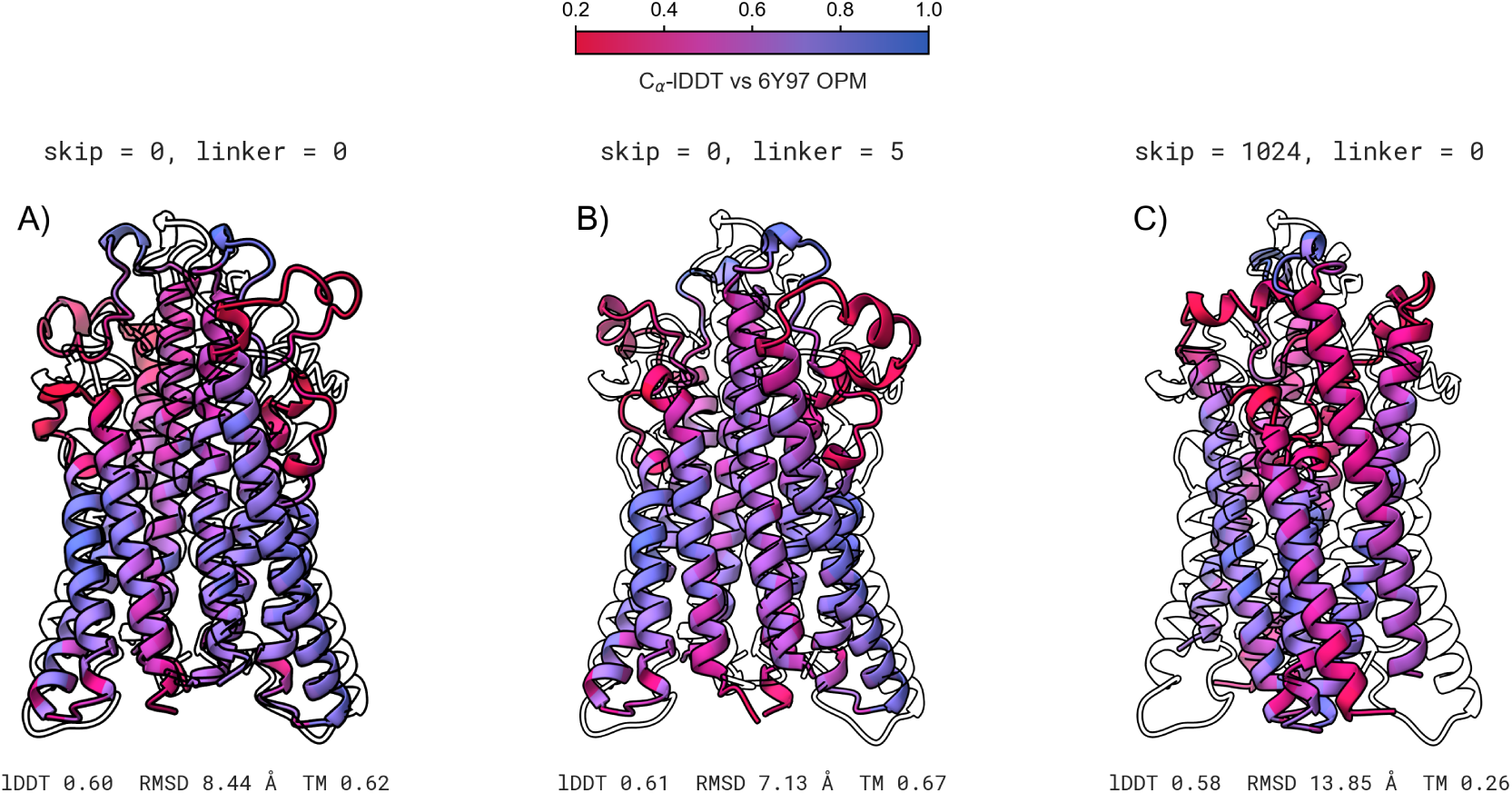
Per-Residue Cα-lDDT of Three ESMFold-Sweep CD20 Dimer Configurations. OPM-aligned ESMFold *apo* CD20 dimer predictions for the two low-RMSD contiguous-encoding cells (`skip = 0, linker = 0 skip = 0, linker = 5`) and the BAGEL recipe (`skip = 1024, linker = 0`) from the Figure 15 matrix. Each cartoon is the prediction superposed on the `6Y97` OPM reference (black silhouette) and colored by per-residue Cα-lDDT [33] (red = poorly preserved local environment, blue = high preservation; inclusion radius 15 Å). In-panel numbers report global Cα-lDDT and Cα RMSD against the OPM reference. The two low-RMSD cells track the OPM ghost across the membrane core, while the BAGEL recipe dimer is visibly displaced.

### S9 BAGEL-CAR proliferation outcome correlations

Using data from the original paper [6], we examine how well proliferation enrichment can be predicted for BAGEL-CAR designs alone, excluding designs from the other teams. We screened all 402 numeric columns in the competition metric table against two outcomes: (i) oriented area under the receiver operating characteristic curve (AUROC) for classifying *enriched* vs *not-enriched* (the latter including the single depleted design) among the 330 detected designs, and (ii) absolute Spearman *ρ* with log fold change (logFC). For each design strategy, we report the best single metric and assess its significance via a best-of-metrics permutation test (200 label shuffles; each shuffle repeats the full 402-metric search, yielding the null distribution of the best achievable score by chance).

Figure 17 shows the AUROC and |*ρ*| distributions across all 402 metrics. We restrict the strategy-level screen to `SOL`, `LNK-BB`, and `LNK-AB` — the three strategies with enough designs in both outcome classes for a meaningful comparison. Among recovered designs, `BGL` has no enriched designs (0/17), whereas all six recovered `LNK-AA` designs are enriched (6/6); because each strategy contains only one outcome class, AUROC is not estimable for either. The strongest AUROC is 0.624 (`esmfold_ipae_end`, permutation *p* = 0.015) and the strongest |*ρ*| is 0.328 (`esmfold_n_hs_engaged_end`, *p* = 0.005). None of the metrics provided by Chai-1 or Boltz-1 appears predictive of outcome, despite both models generally placing binders on the extracellular side more oŁen than ESMFold, as shown in Figure 4. While the best metrics reach nominal significance (AUROC *p* = 0.015, |*ρ*| *p* = 0.005), no individual per-strategy metric shows consistent predictive power, so no single metric emerges as a reliable predictor of outcome for BAGEL-CAR designs.

**Figure 17:**
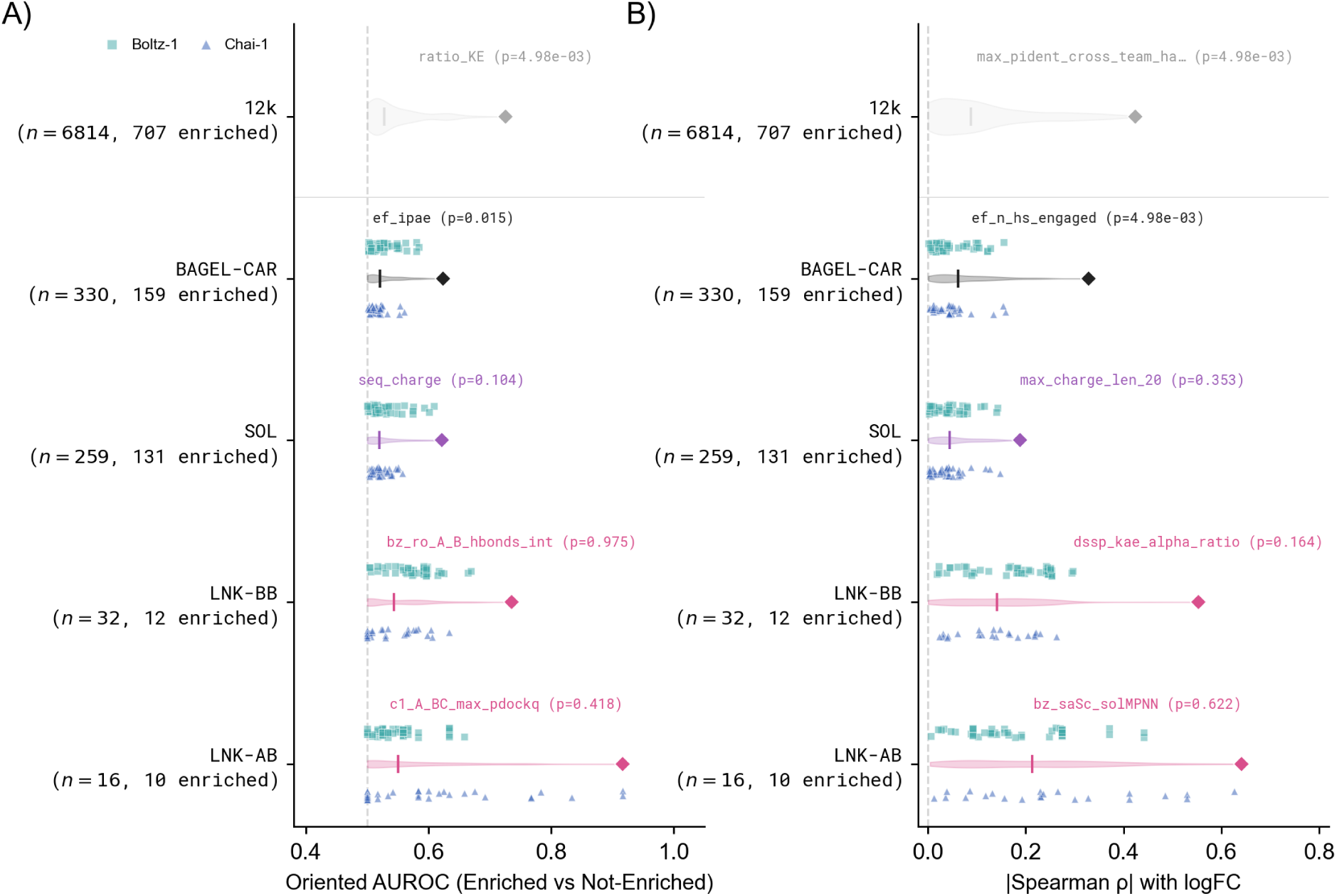
BAGEL-CAR Metric-vs-Outcome Correlations. Distribution of **A)** oriented AUROC (enriched vs not-enriched) and **B)** absolute Spearman *ρ* with logFC across all 402 competition-analyzed metrics (as provided in Kosonocky et al. [6]), for all BAGEL-CAR designs and for each strategy. Violins show the per-group distribution (width scaled by *n*); vertical ticks mark the median; colored diamonds mark the best single metric. Annotations give the best metric name (abbreviated) and best-of-metrics permutation *p*-value (200 permutations). Individual Boltz-1 metrics (teal squares, *n* = 49) and Chai-1 metrics (blue triangles, *n* = 40) are overlaid above and below each violin, respectively, showing that structure-prediction confidence scores scatter across the full distribution without clustering at the predictive end. The dashed line marks chance level (AUROC = 0.5 in A; |*ρ*| = 0 in B). The 12k row at the top shows the competition-wide screen as reference [6]. `BGL` (0 enriched designs) and `LNK-AA` (0 not-enriched among recovered) are omitted because AUROC is not estimable for either.

### S10 Representative structure predictions

**Figure 18:**
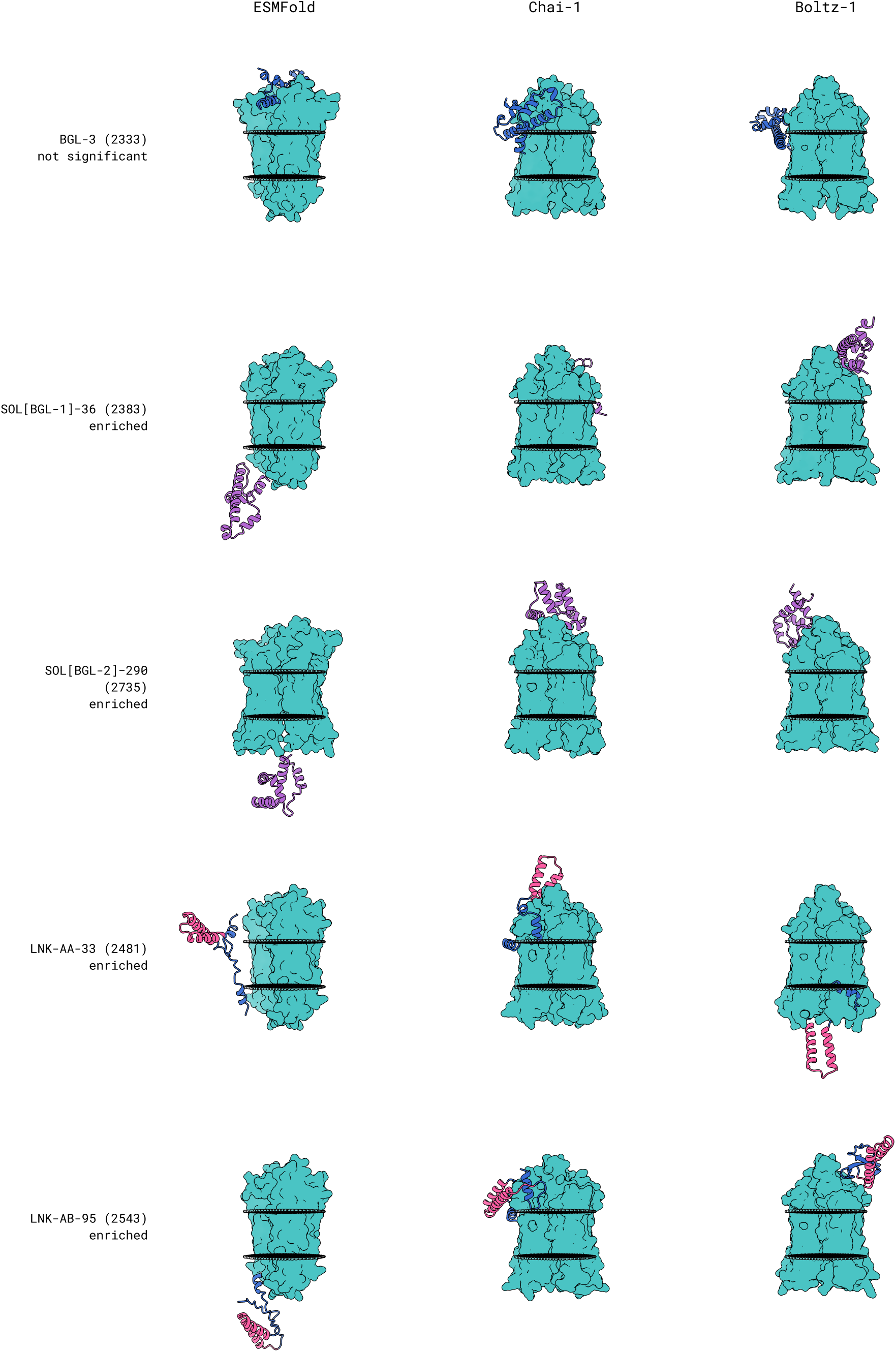
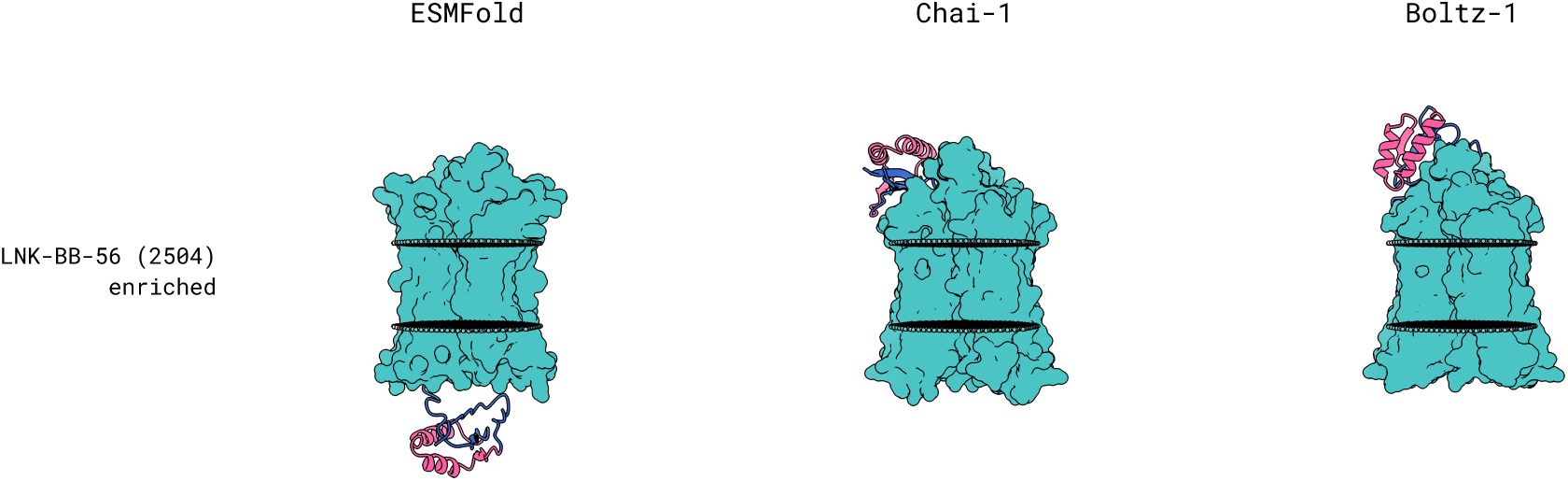
Representative BAGEL-CAR Structure Predictions from ESMFold, Chai-1, and Boltz-1. Predicted CD20 dimer as teal surface; binder as cartoon. `BGL` binders blue, `SOL` binders purple; `LNK` binders show BAGEL-derived termini in blue and the RFdiffusion linker in pink. ESMFold panels use the BAGEL-chain-order `CD20A:CD20B:binder` encoding with `position_ids_skip=1024` and no explicit glycine linker. Structures are aligned to the OPM-oriented CD20 dimer (`6Y97`). Faint gray spheres mark the OPM membrane boundary (the `6Y97`-OPM dummy-atom planes); each structure is shown in its OPM membrane orientation, with the extracellular side up and the intracellular side down. Row labels report competition pooled-screen outcomes: `enriched` = FDR-significant ≥ 2-fold differential expansion in CD20-positive Raji co-culture versus the no-target control; `not significant` = detected but without significant differential expansion. Design `SOL[BGL-1]-36` (`2383`) is the top validated BAGEL-CAR competition hit.

